# Sensory Modality and Spoken Language Shape Reading Network in Blind Readers of Braille

**DOI:** 10.1101/2021.08.24.457544

**Authors:** Mengyu Tian, Elizabeth J. Saccone, Judy S. Kim, Shipra Kanjlia, Marina Bedny

## Abstract

The neural basis of reading is highly consistent across many languages and scripts. Are there alternative neural routes to reading? How does the sensory modality of symbols (tactile vs. visual) influence their neural representations? We examined these questions by comparing reading of visual print (sighted group, n=19) and tactile Braille (congenitally blind group, n=19). Blind and sighted readers were presented with written (words, consonant strings, non-letter shapes) and spoken stimuli (words, backward speech) that varied in word-likeness. Consistent with prior work, the ventral occipitotemporal cortex (vOTC) was active during Braille and visual reading. A posterior/anterior vOTC word-form gradient was observed only in sighted readers with more anterior regions preferring larger orthographic units (words). No such gradient was observed in blind readers. Consistent with connectivity predictions, in blind compared to sighted readers, posterior parietal cortices were recruited to a greater degree and contained word-preferring patches. Lateralization of Braille in blind readers was predicted by laterality of spoken language and reading hand. The effect of spoken language increased along a cortical hierarchy, whereas effect of reading-hand waned. These results suggested that the neural basis of reading is influenced by symbol modality and spoken language and support connectivity-based views of cortical function.

## Introduction

Written language is among the most impressive human cultural achievements. The capacity to record and transmit information over space and time has enabled the accumulation of scientific, technological, and historical knowledge across generations and continents. How does the human brain accommodate this cultural invention, which emerged only approximately 5,000 years ago?

Despite being a recent cultural invention, the neural basis of reading is highly consistent across a variety of languages and visual scripts, including alphabetic, logographic (e.g., Chinese), and syllabic writing systems (e.g., Japanese Kana) (Bolger et al. 2005; Hu et al. 2010; Nakamura et al. 2012; Rueckl et al. 2015; Krafnick et al. 2016; Feng et al. 2020; Tian et al. 2020). All of these reading systems engage regions within the left lateral ventral occipitotemporal cortex (vOTC). A region in the left lateral vOTC has been termed the ‘visual word form area’ (VWFA) because of its preferential response to written words and letter combinations over non-linguistic visual stimuli and speech (Cohen et al. 2000; Baker et al. 2007; Dehaene et al. 2010; Dehaene and Cohen 2011). The VWFA is situated within a posterior/anterior processing gradient. During reading, visual symbols are first processed by early visual cortices and posterior portions of vOTC, which represent simple visual features (e.g., line junctions) (Dehaene et al. 2005; DiCarlo and Cox 2007). By contrast, the middle and anterior potions of lateral vOTC are specialized for progressively larger orthographic units, from written letters, letter combinations/bigrams, and finally whole words (Dehaene et al. 2005; Binder et al. 2006; Vinckier et al. 2007).

An open question is whether the vOTC’s posterior/anterior processing stream is the only way for the brain to implement reading and, relatedly, why the neural basis of reading takes this particular form. Comparing tactile Braille reading among congenitally blind individuals to print reading in sighted people offers unique insights into the causal mechanisms that determine the neural basis of reading. Braille is read by passing the fingers along raised dot patterns, with each Braille character a three-rows-by-two- columns dot matrix (Millar 2003). This distinctive tactile reading system provides insight into whether and how the sensory modality of symbols influences their neural representations. Do blind readers of Braille likewise show an anterior/posterior orthographic gradient in the vOTC?

A prominent view holds that, past the initial sensory entry points in V1 (print) and S1 (Braille), reading depends on the same neural mechanisms in blind Braille readers and sighted print readers (Büchel et al. 1998; Reich et al. 2011; Debowska et al. 2016; Rączy et al. 2019). This view is based on the observation that reading Braille elicits activation in the anatomical location of the ‘VWFA’ (Reich et al. 2011; Debowska et al. 2016; Siuda-Krzywicka et al. 2016; Bola et al. 2019; Rączy et al. 2019; Dzięgiel- Fivet et al. 2021). At the same time, no prior study to our knowledge has tested for the presence of an orthographic gradient in vOTC of blind readers. The available work has only looked at whether the anatomical location of the VWFA responds to Braille. Unlike visual print, Braille does not enter the vOTC from V1, but rather originates in primary somatosensory cortex (S1). Does a posterior/anterior orthographic gradient emerge despite this difference in neural entry points?

The first goal of the current study was to test whether sighted print and blind Braille readers recruit a similar posterior/anterior orthographic gradient within vOTC. To answer this question, we compared responses during Braille (blind) and visual print (sighted) reading of analogous written stimuli of different orthographic richness (words, consonant strings, and unfamiliar shapes (control)). The same participants were also presented with spoken words and backward speech control stimuli, to enable comparison of written and spoken language processing.

An alternative, though not a mutually exclusive possibility, is that regions closer to somatosensory cortices, in parietal cortex, play a special role in Braille reading. In sighted readers, the vOTC is the culmination of the visual object recognition stream and occupies a key connectivity position between visual input on the one hand and linguistic representations on the other (Yeatman et al. 2013; Bouhali et al. 2014; Hannagan et al. 2015; Saygin et al. 2016; Stevens et al. 2017; Barttfeld et al. 2018; Li et al. 2020). In the case of tactile reading, posterior parietal cortices (PPC) arguably occupy an analogous connectivity-based position for blind readers of Braille. The PPC lies adjacent and posterior to early somatosensory cortex (SMC) and plays a key role in tactile shape and texture perception and higher order tactile processing (Hegner et al. 2010; Bauer et al. 2015). Like the ‘VWFA’, PPC is anatomically connected to language and working memory systems (Duhamel et al. 1998; Lewis and Van Essen 2000; Kaas 2012; Ruschel et al. 2014; Burks et al. 2017). Whether the PPC of proficient blind Braille readers contains Braille specialization, akin to specialization for visual letters and words found in vOTC of sighted readers, has not yet been systematically investigated.

Several previous neuroimaging studies have documented PPC activity during Braille reading tasks but have not explored whether these responses are Braille-specific or distinct from what is observed in sighted readers in analogous tasks (Sadato et al. 1998; Burton, Snyder, Conturo, et al. 2002; Burton et al. 2012; Siuda-Krzywicka et al. 2016; Dzięgiel-Fivet et al. 2021). Previous studies also find that early somatosensory cortices (SMC) show expanded finger representations in proficient Braille readers, however, preferences for Braille over matched non-Braille tactile stimuli have not been found in SMC (Pascual-Leone and Torres 1993; Pascual-Leone et al. 1993; Sadato et al. 1998; Burton, Snyder, Conturo, et al. 2002; Burton et al. 2004; Kupers et al. 2007). Thus, it is unknown if any cortical areas outside of vOTC acquires specialization for Braille letters and words in blind readers.

We used sensitive region of interest (ROI) analyses as well as complementary data-driven cortical gradient maps to test whether regions within PPC show preferential responses to Braille over non-linguistic but perceptually similar tactile stimuli on the one hand and preferential responses to Braille over spoken language on the other, akin to the functional profile of the ‘VWFA’ in sighted readers. We hypothesized that analogous to the vOTC gradient, word-preferring portions of PPC should be found further from, and consequently posterior to, primary somatosensory cortices. Such analysis approaches are particularly relevant in the context of understanding the neural basis of reading, since previous studies find that in sighted readers preferential responses to print in vOTC are ‘islands’ among swaths of cortex that respond to complex visual shapes (Cohen and Dehaene 2004; Hannagan et al. 2015; Behrmann and Plaut 2020). Conventional whole-brain analyses that average across participants might therefore miss Braille specialization in PPC.

Finally, our third goal was to test the hypothesis that neural localization of spoken language influences the localization of Braille, analogous to what is observed in readers of visual print. In sighted readers, strong connectivity to spoken language networks predicts the localization of reading networks within vOTC and the lateralization of written language follows that of spoken language (Saygin et al. 2016; Stevens et al. 2017; Li et al. 2020). Reading, like spoken language, is on average strongly left-lateralized in sighted people but those sighted readers whose spoken language responses are localized to the right hemisphere also show right-lateralized written language (Schlaggar and McCandliss 2007; Cai et al. 2010; Seghier and Price 2011; Van der Haegen et al. 2012; Ossowski and Behrmann 2015). It is unknown whether reading and spoken language networks co-lateralize in blind readers as they do in sighted people.

Unlike the strong left-lateralization of spoken language in sighted people, lateralization of spoken language is highly variable across congenitally blind individuals and on average only weakly left-lateralized (Röder et al. 2000, 2002; Lane et al. 2017). This variability enabled us to use individual difference analyses to test whether lateralization of Braille follows that of spoken language across blind individuals. We also tested the effect of reading hand on lateralization and predicted that cortical areas situated at earlier stages of Braille recognition (i.e., S1) would show stronger effects of reading hand, whereas the effect of spoken language laterality would emerge in orthographic and higher-order language regions (PPC, vOTC and inferior frontal cortex).

## Method

### Participants

Nineteen congenitally blind (12 females, mean age = 39.05 years, SD = 13.12) and 19 sighted controls (11 females, mean age = 36.36 years, SD = 19.45) participated in the current study (Table 1). All participants were native English speakers, and none had suffered from any known cognitive or neurological disabilities (screened through self-report). All of the blind participants reported at least some college education, with most having completed a college degree. The blind and sighted groups were matched on age (*t* _(36)_ = 0.499, *p* = 0.621) and years of education (*t* _(36)_ = 0.867, *p* = 0.392). Blind participants had at most minimal light perception from birth. Blindness was caused by pathology anterior to the optic chiasm (i.e., not due to brain damage). All blind participants were fluent, frequent Braille readers who began learning Braille at an average age of 4.6 years (SD = 1.53), rated their reading ability as proficient to expert (mean = 4.6, SD =0.69 on a scale of 1 to 5), and reported reading Braille for an average of 19.94 hours per week. We obtained information on Braille-reading hand dominance through a post-experimental survey conducted over the telephone with 17 of the 19 blind adult participants (Table 1). All participants gave informed consent according to procedures approved by the Johns Hopkins Institutional Review Board.

**Table 1.** Participant information

| Participant no. | Age (y) | Gender | Handedness | Reading handedness | Levels of education | Education years | Cause of blindness | Age started reading Braille (y) | Reading hours per week | Self-reported Braille reading ability (1-5) |
| --- | --- | --- | --- | --- | --- | --- | --- | --- | --- | --- |
| B1 | 21 | M | L | Bi-R | SC | 16 | LCA | 4 | 14 | 5 |
| B2 | 64 | F | R | Bi-R | BA | 16 | ROP | 6 | 56 | 5 |
| B3 | 53 | M | R | Bi-R | JD | 20 | LCA | 6 | 7 | 4 |
| B4 | 34 | M | R | L | SC | 15 | Born without optic nerve | 3 | 21 | 5 |
| B5 | 42 | M | Am | L | BA | 17 | ROP | 3 | 21 | 5 |
| B6 | 29 | M | R | Bi-L | SC | 14 | LCA | 4 | <1 | 4 |
| B7 | 39 | F | R | L | BA | 16 | ROP | 4 | 2 | 5 |
| B8 | 34 | F | R | -- | SC | 15 | Optic Nerve Detached | 3 | -- | 5 |
| B9 | 49 | M | R | Bi-R | BA | 17 | unknown | 8 | <1 | 3 |
| B10 | 26 | F | R | Bi-R | MA | 19 | ROP | 3 | 56 | 3 |
| B11 | 49 | F | L | R | MA | 17 | LCA | 7 | 14 | 5 |
| B12 | 39 | F | R | L | MA | 21 | ROP | 5 | 14 | 5 |
| B13 | 35 | F | R | Bi-L | MA | 19 | LCA | 4 | 14 | 5 |
| B14 | 46 | F | R | -- | BA | 17 | ROP | 4 | -- | 5 |
| B15 | 33 | F | R | L | BA | 17 | ROP | 4 | 14 | 4 |
| B16 | 25 | F | Am | Bi-R | MA | 19 | LCA | 5 | 56 | 5 |
| B17 | 23 | M | R | Bi-R | BA | 15 | LCA | 4 | 28 | 5 |
| B18 | 33 | F | A | R | BA | 16 | LCA | 3 | 14 | 5 |
| B19 | 68 | F | R | Bi-R | MA | 19 | ROP | 5 | 7 | 5 |
| **Average** |  |  |  |  |  |  |  |  |  |  |
| Blind (n=19) | 39.05 (SD=13.12) | 12F | 2L/3Am | -- | BA | 17.11 (SD = 1.91) | -- | 4.6 (SD=1.53) | 19.94 (SD=18.36) | 4.6 (SD=0.69) |
| Sighted (n=19) | 36.36 (SD=19.45) | 11F | 2 L | -- | BA | 16.53 (SD = 2.19) | -- | -- |  | -- |

Handedness: left (L), ambidextrous (Am), or right (R), based on Edinburgh Handedness
Inventory. BA = Bachelor of Arts; MA = Master of Arts; HS = High School; JD = Juris Doctor; SC
= Some College; ROP = Retinopathy of prematurity; LCA = Leber's congenital amaurosis. For
Braille ability, participants were asked: "On a scale of 1 to 5, how well are you able to read Braille,
where 1 is 'not at all', 2 is 'very little', 3 is 'reasonably well', 4 is 'proficiently', and 5 is 'expert'?"

### Stimuli and experimental procedures

The fMRI task included three reading conditions: words, consonant strings, and non-letter shapes (Figure 1) and two listening conditions: words, backwards speech (Kim et al. 2017). Reading stimuli were visual for sighted participants and tactile for blind participants. Braille words were written in Grade-II contracted English Braille, which is the most common form of Braille in the United States and therefore the most ecologically valid form of written material for this population. Visual words were matched to Braille words on average character length. Each consonant string stimulus consisted of 4 visual/Braille consonants. The tactile control stimuli consisted of 24 unique strings of 4 non-letter shapes made of raised Braille dots. The shapes were chosen based on pilot testing with a blind proficient Braille reader who reported them to be recognizable and not confusable with Braille letters. The piloting process revealed that it was critical to create shapes that did not fit into a single Braille cell (2 x 3 array of pins), otherwise the shapes would be temporarily confused with Braille letters and characters. We therefore created shapes/pin arrays ranging in size from 4 × 5 to 7 × 7 pins. The visual control stimuli were 24 unique strings, each comprised of 4 characters, which were false fonts. These were matched to English consonants on the number of strokes, ascenders, descenders and stroke thickness (Kim et al. 2017). The auditory words were recorded by a female native English speaker. The average word length was 5 letters long (SD = 1.4 letters) and the average playtime was 0.41 s (SD = 0.3 s). The control auditory stimuli comprised backward speech sounds created by playing each audio word in reverse.

**Figure 1.**
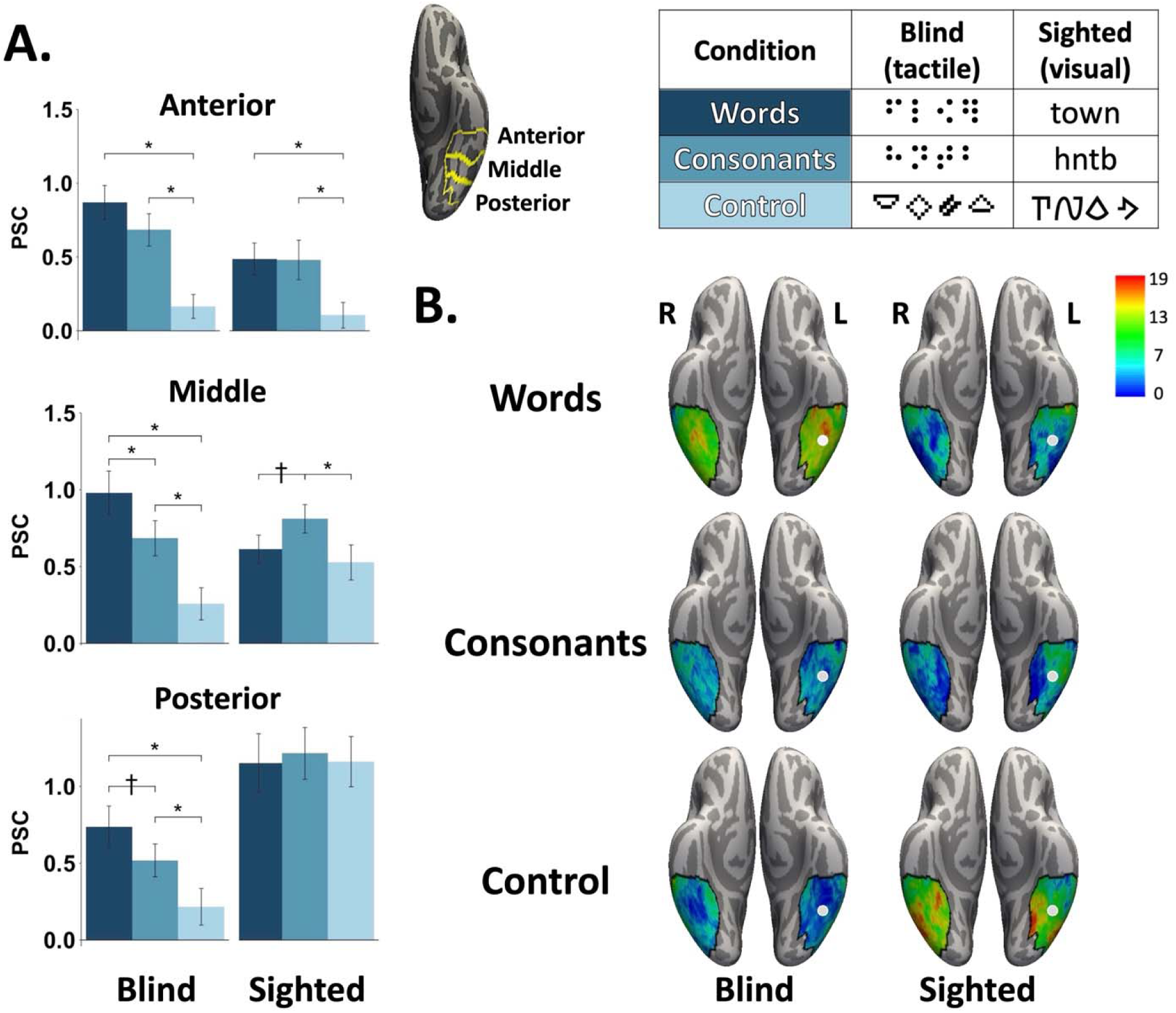
(A) Responses in left vOTC across the posterior, middle, and anterior subregions for blind and sighted groups during the reading tasks. Bars show results from consonant string > control shapes leave-one-run out individual subject ROI analysis. Error bars denote standard errors +/- the mean. Asterisks (*) denote significant Bonferroni-corrected pairwise comparisons (*p* < 0.05). (B) Group histogram plots for topographical preferences map. Each vertex was color-coded according to the number of participants showing preference for words, consonants, or control shapes. Silver circles mark previously reported location of VWFA (MNI coordinate: -46, -53, -20)(McCandliss et al. 2003).

On each trial, participants were presented with 6 stimuli from a single condition (e.g., Braille words) followed by a memory probe. Participants judged whether the probe had appeared among the previous 6 stimuli. This task was used to ensure attentive processing. The experiment had a total of 5 runs, each with 20 trials. The blind participants were asked to read with their dominant hand and responded with the other hand. Each condition was repeated 4 times per run, and the order of conditions was counterbalanced across runs. There were 6 rest periods (16 s) throughout each run. One sighted participant and two blind participants were excluded from behavioral analysis due to failure to record their responses (see SI and Kim et al., 2017 for details).

### fMRI data acquisition

Functional and structural images were acquired using a 3T Phillips scanner at the F. M. Kirby Research Center. T1-weighted images were collected using a magnetization-prepared rapid gradient-echo (MP- RAGE) in 150 axial slices with 1 mm isotropic voxels. Functional BOLD scans were collected T1-weighted structural images were collected in 150 axial slices with 1 mm isotropic voxels. Functional BOLD scans were collected in 36 sequential ascending axial slices. TR = 2 s, TE = 30 ms, flip angle = 70°, voxel size = 2.4 × 2.4 × 2.5 mm, inter-slice gap = 0.5 mm, field of view (FOV) = 192 × 172.8 × 107.5.

### fMRI data analysis

#### Preprocessing and whole-cortex analysis

Analyses were performed using FSL (version 5.0.9), FreeSurfer (version 5.3.0), the Human Connectome Project workbench (version 1.2.0), and custom in-house software. The cortical surface was created for each participant using the standard FreeSurfer pipeline (Dale et al. 1999; Smith et al. 2004; Glasser et al. 2013). For task data, preprocessing of functional data included motion-correction, high-pass filtering (128 s cut-off), and resampling to the cortical surface. Cerebellar and subcortical structures were excluded. On the surface, the task data were smoothed with a 6 mm FWHM Gaussian kernel.

Three conditions of the reading task and two conditions of the listening task were included in a general linear model. Analysis focused on the time-period during the initial six stimuli of each trial. Probe stimulus and response periods were modeled separately and are not reported. White matter signal, CSF signal, as well as motion spikes, were included as the covariates of no interest.

Whole-cortex random-effects analysis were run using mixed-effects and thresholded at *p* < 0.01 vertex-wise, and *p* < 0.05 cluster-wise, Family Wise Error corrected for multiple comparisons across the cortex.

#### fMRI ROI analysis

We used subject-specific functional ROI analyses to examine five regions of interest: vOTC, V1, inferior frontal cortex (IFC), and posterior parietal cortex (PPC), and the hand region of the left primary somatosensory-motor cortex (SMC) (see SI for the search space construction details). Individual-subject functional ROIs were defined within each of the above search spaces using a leave-one-run-out cross-validation procedure (e.g., Ellis et al., 2021; Kim et al., 2017; Saygin et al., 2016). This procedure ensures that independent sets of data are used to define ROIs and to test hypotheses and effectively uses each run as a ‘localizer scan’ for the other runs of data (Cohen et al., 2019; Ellis et al., 2021; Kanjlia et al., 2016, 2021; Ratan Murty et al., 2020; Saygin et al., 2016). ROIs were defined based on data from all but one run, then the percent signal change (PSC) was extracted from the left-out run. This procedure was repeated iteratively across all runs and the PSC was averaged across iterations (see SI for details). Each individual subject’s ROI was defined as the top 5% of vertices activated for tactile/visual consonant strings > tactile/visual shapes. The main text reports results for ROIs defined using the consonant string > control contrasts so as to focus on orthographic as opposed to semantic responses. In the supplement we report results for ROIs defined using the words > control contrasts, which produced highly similar results (see Figure S3 and Figure S6). Repeated-measured ANOVAs were used to analyze the ROI data, and two-tailed paired *t*-tests were used for pairwise comparisons. All *p* values were Bonferroni-corrected (see SI for details).

A vector of ROI analysis were conducted in PPC mask (Konkle and Caramazza 2013). We divided the PPC mask into 13 segments centered along an equal distanced spline and plotted average activity for three tactile conditions in each segment (see SI for details).

#### Topographical preference map

To examine topographic gradients in bilateral vOTC and PPC with data-driven winner-take-all approaches. Each vertex was color-coded according to which stimulus condition showed highest activity. We also calculated the topographical preferences winner-take-all map for each participant, then created group histogram plots where each vertex was color-coded according to the number of participants showing preference for a certain condition (i.e., words, consonant strings or control).

#### Laterality index analysis

LI was calculated separately for the reading and listening tasks for each participant in the SMC, PPC, vOTC, and IFC. For the reading task, LI was determined based on the tactile/visual words > rest contrast. For the listening task, LI was determined using the audio words > rest contrast. The LI was calculated using the standard formula: (L - R) / (L + R), where L and R refer to the sums of the *z* statistics from the relevant contrast within the left and right hemispheres, respectively. LI ranges from -1 to 1, with a score of 1 indicating strong left lateralization and -1 strong right lateralization. The bootstrap/histogram method was used to ensure that LIs were not overly influenced by arbitrary activation threshold choices or outlier voxels (Wilke and Schmithorst 2006). Bootstrapped LIs were computed using 20 evenly spaced thresholds ranging from *z* = 1.28 to *z* = 4.26 (corresponding to one-sided *p* = 0.1 to *p* = 0.00001, uncorrected). The LI reported for each participant represents the average across all thresholds. Participants were excluded from the LI analysis if they did not have suprathreshold activation in both hemispheres (see SI for details).

## Results

### Behavioral Results

Sighted and blind readers alike were more accurate at remembering more word-like stimuli in both the reading and listening tasks (reading: main effect of lexicality (words and consonant strings > control shapes): *F*_(2, 64)_ = 24.991, *p* < 0.001; listening (words > control shapes): *F*_(1, 32)_ = 46.137, *p* < 0.001. That is, both groups achieved higher accuracy rates when reading words (sighted group visual words: 91.2%, blind group tactile words: 83.1%) and consonant strings (sighted group: 86.6%, blind group: 77.9%), than control shapes strings (sighted group: 72.7%, blind group: 62.5%). Likewise, accuracy rates were higher for audio words (sighted group: 91.3%, blind group: 88.4%) than backwards speech (sighted group: 75.9%, blind group: 64.2%) (see Figure S1). There was no lexicality by group interaction in either reading (*F*_(2, 64)_ = 0.067, *p* = 0.935) or listening tasks (*F*_(1, 32)_ = 2.252, *p* = 0.143). Sighted readers had slightly higher accuracy rates on both reading and listening tasks compared to blind readers (reading main effect of group: *F*_(1, 32)_ = 8.21, *p* < 0.01; listening main effect of group: *F*_(1, 32)_ = 6.305, *p* < 0.05).

Participants were also faster at responding to word-like stimuli across reading (words and consonant strings < control shapes; *F*_(2, 64)_ = 12.998, *p* < 0.001) and listening tasks (words < control shapes; *F*_(1, 32)_ = 30.763, *p* < 0.001). Sighted readers were faster than blind readers for the reading task (main effect of group: *F*_(1, 32)_ = 10.641, *p* < 0.01), but not for the listening task (main effect of group: *F*_(1, 32)_ = 2.488, *p* = 0.125). For the reading task, there was a group by lexicality interaction (*F*_(2, 64)_ = 3.943, *p* < 0.05). Pairwise comparisons showed shorter reaction times for words and consonant strings relative to the control condition in the blind group (words vs. control shapes, *t*_(32)_ = -4.208, *p* < 0.01; consonant strings vs. control shapes, *t*_(32)_ = -3.722, *p* < 0.05; words vs. consonant strings, *t*_(32)_ = -1.177, *p* = 0.24). For the sighted group, the only difference was faster reaction times for words than consonant strings (words vs. control shapes, *t*_(32)_ = -1.573, *p* =0.377; consonant strings vs. control shapes, *t*_(32)_ = -0.718, *p* > 0.99; words vs. consonant strings, *t*_(32)_ = -3.084, *p* < 0.05; see Figure S1).

### fMRI Results

#### Visual (sighted) but not tactile Braille reading (blind) elicits a posterior-to-anterior functional gradient in left vOTC and shows left-lateralization

We divided the left and right vOTC into posterior, middle, and anterior subregions (ROIs) and observed a different posterior-to-anterior shift and a different lateralization pattern across visual and Braille reading (Figure 1A). A four-way hemisphere (left, right) by posterior/anterior subregion (posterior, middle, anterior) by lexicality (words, consonant strings, control shapes) by group (sighted, blind) ANOVA on the reading task revealed a four-way interaction (*F* (4, 144) = 2.954, *p* < 0.05), indicating that lexicality, hemisphere, and posterior/anterior subregion interacts with group (see SI for a complete summary of all effects). As predicted, the sighted group showed the previously documented posterior-to-anterior functional gradient in left but not right vOTC: larger responses to words in anterior vOTC, to consonant strings in middle vOTC and no differences between visual conditions in posterior vOTC (three-way interaction between hemisphere (left, right), posterior/anterior subregion (posterior, middle, anterior) and lexicality (words, consonant strings, control shapes): *F* _(4, 72)_ = 4.344, *p* < 0.01; in left vOTC two-way interaction between lexicality (words, consonant strings, control shapes) and posterior/anterior subregion (posterior, middle, anterior): *F* _(4, 72)_ = 8.237, *p* < 0.001) (see SI for pairwise comparisons within each subregion). By contrast, no gradient or laterality differences were observed in the blind group. Rather, all subregions of bilateral vOTC responded most to words, followed by consonant strings followed by control shapes (Figure 1A) (three-way hemisphere (left, right) by posterior/anterior subregion (posterior, middle, anterior) by lexicality (words, consonant strings, control shapes) ANOVA interaction: *F* _(4, 72)_ = 0.747, *p* = 0.563) (see SI for pairwise comparisons and listening task results). A similar word preference was observed in V1 for blind readers (Sadato et al. 1996; Cohen et al. 1997)(see SI for details).

The group differences identified by ROI analyses (above) were confirmed by data-driven topographical preference maps where each vertex is color-coded according to the number of participants showing numerical preference for a particular condition (Figure 1B, also see SI Figure S10 for topographical preference maps in each participant and Figure S4 for topographical preference maps in group average). In the sighted group, the word-preferring vertices were concentrated most consistently across participants in the left anterior vOTC. The location of the consonant-preferring vertices was more distributed, but the left anterior and middle vOTC were still the areas with the largest concentration of subjects. The location of the control shapes-preferring vertices were most often observed in left posterior and medial vOTC in the sighted. In the blind group, most participants showed a preference for words throughout the vOTC. The highest degree of cross-participant overlap was observed bilaterally at MNI coordinates (-44, -43, -18) and (41, -48, -22).

#### The posterior parietal cortex (PPC) but not S1 of blind readers shows a preference for written Braille words and consonant strings

Consistent with the hypothesis of Braille specialization in PPC of congenitally blind readers, the PPC of blind readers preferred tactile over auditory stimuli and among tactile stimuli it preferred Braille words and consonant strings over tactile control stimuli. The two-way lexicality (words, consonant strings, control shapes) by group (sighted, blind) ANOVA revealed a marginally different response profile across blind Braille and sighted print readers in the reading task (lexicality by group interaction effect: *F* _(2, 72)_ = 2.682, *p* = 0.075; main effect of lexicality: *F* _(2, 72)_ = 12.482, *p* < 0.001; main effect of group: *F* _(1, 36)_ =0.081, *p* = 0.778).

In blind readers, the PPC responded more to Braille words and Braille consonant strings than control shapes (words vs. control shapes: *t*_(36)_ = 3.602, *p* < 0.05; consonant strings vs. control shapes: *t*_(36)_ = 3.595, *p* < 0.05; words vs. consonant strings: *t*_(36)_ = 1.288, *p* > 0.99; Bonferroni-corrected paired *t*-tests) (Figure S5). In the sighted group, words were not different from control shapes but consonant strings elicited higher responses than control shapes, and words and consonants did not differ from each other (words vs. control shapes: *t*_(36)_ = 1.161, *p* > 0.99; consonant strings vs. control shapes: *t*_(36)_ = 3.282, *p* < 0.05; words vs. consonant strings: *t*_(36)_ = -2.208, *p* = 0.75; Bonferroni-corrected paired *t*-tests).

The PPC also showed larger responses to tactile than auditory stimuli in the blind group (two-way modality (tactile, auditory) by lexicality (word, control) ANOVA, main effect of modality: *F* _(1, 18)_ = 11.919, *p* < 0.01; main effect of lexicality: *F* _(1, 18)_ =9.079, *p* < 0.01; modality by lexicality interaction: *F* _(1, 18)_ = 4.414, *p* = 0.05). Responses to visual and auditory stimuli in the sighted were not different (two-way modality (visual, auditory) by lexicality (word, control) ANOVA, main effect of modality: *F* _(1, 18)_ =0.478, *p* = 0.498; main effect of lexicality: *F* _(1, 18)_ = 3.762, *p* = 0.068; modality by lexicality interaction: *F* _(1, 18)_ =2.308, *p* = 0.146).

There was a larger response to auditory words than auditory control stimuli in both groups in the PPC (two-way lexicality (audio words, audio control) by group (sighted, blind) ANOVA, main effect of lexicality: words > control, *F* _(1, 36)_ = 7.624, *p* < 0.01; main effect of group: *F* _(1, 36)_ =0.602, *p* = 0.443; group by lexicality interaction: *F* _(1, 32)_ = 2.493, *p* = 0.123, Figure S5).

A data-driven topographic winner-take-all map showed that preferential responses to Braille words were located in the most posterior portion of PPC and extended into parieto-occipital and dorsal occipital regions (Figure 2A). A vector-of-ROIs analysis along the anterior/posterior extent of PPC revealed a position by reading condition interaction in blind readers (left PPC: *F* _(12, 24)_ = 16.62, *p* < 0.001; right PPC: *F* _(14, 28)_ = 13.15, *p* < 0.001; see Figure S7 for right PPC). The most anterior portions of PPC, immediately adjacent to S1, showed preferential responses to tactile shapes, whereas posterior PPC showed a preference for Braille words. In the winner-take-all map, a small middle region in left and right PPC showed the highest responses to Braille consonant strings. These results were confirmed by a participant-number-histogram (Figure 2B). This pattern is suggestive of an anterior-to-posterior decoding gradient in PPC in blind Braille readers analogous to the posterior-to-anterior gradient observed in vOTC of sighted print readers.

**Figure 2.**
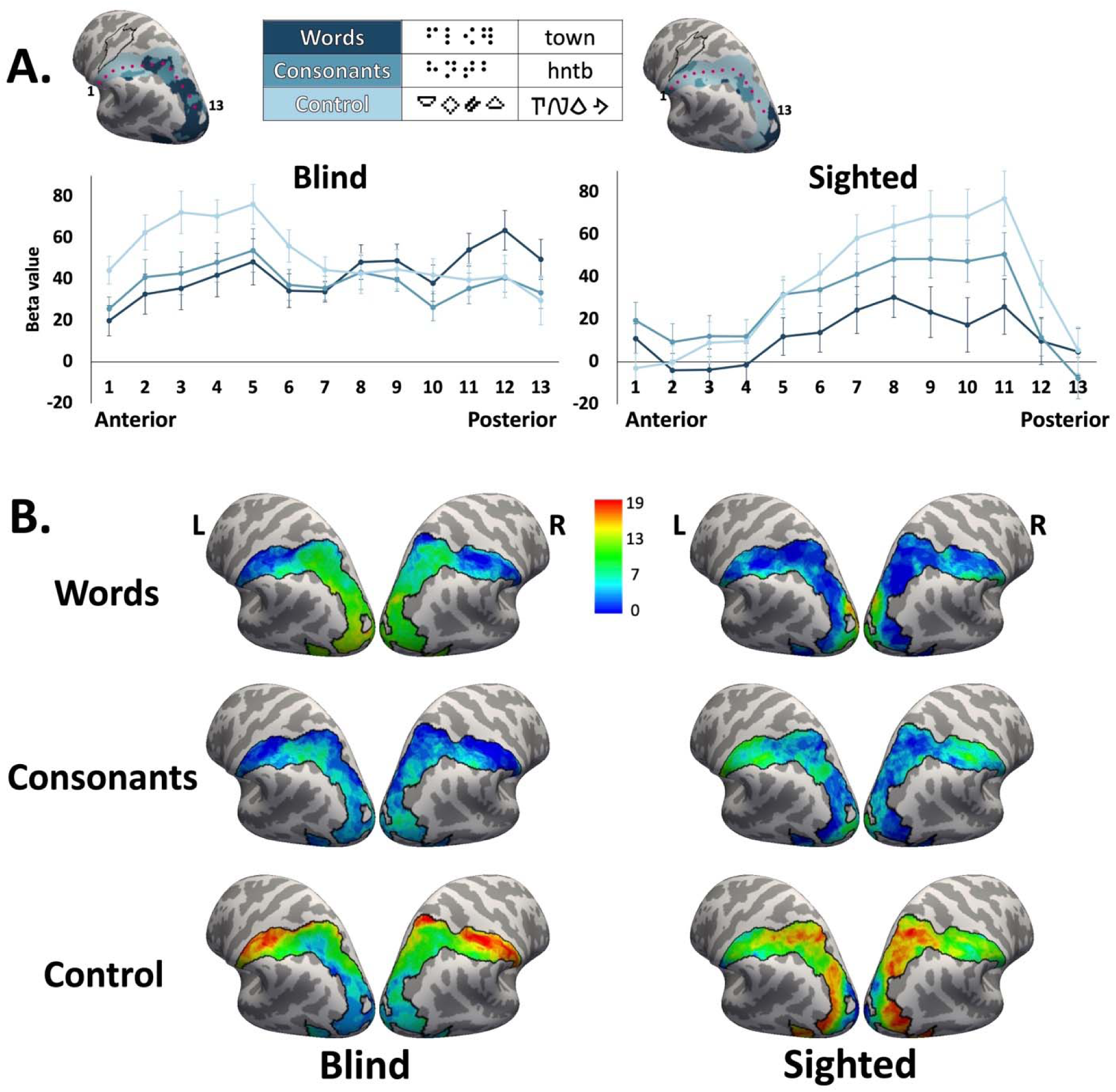
(A) Posterior parietal cortex (PPC) winner-take-all map (upper) during the reading task: words, consonant strings, and control stimuli. The black outline indicates the hand region of the primary sensory-motor cortex (Neurosynth). Vector-of-ROIs results (lower) showed the mean response to each reading condition along left anterior/posterior PPC extent in the blind group (13 segments). Segment centers are marked by red dots (see Figure S7 for right PPC). (B) Group participant-number-histogram plots for topographical preferences winner-take-all map. Each vertex was color-coded according to the number of participants showing preference for words, consonants, or control shapes.

By contrast to the PPC, early somatosensory cortices (SMC hand region, Figure S5) did not show a preferential response to Braille words and showed the same functional profile across blind and sighted readers (two-way lexicality (words, consonant strings, control shapes) by group (sighted, blind) ANOVA, main effect of lexicality: *F* _(2, 72)_ = 7.946, *p* < 0.001; main effect of group: *F* _(1, 36)_ = 2.776, *p* = 0.104; group by condition interaction: *F* _(2, 72)_ = 1.922, *p* = 0.156). (See SI for details of listening task results.) Likewise, the left IFC, a high-level language region, showed a similarly preferential response to linguistic stimuli (i.e., words) across groups and tasks (reading task, two-way lexicality (words, consonant strings, control shapes) by group (sighted, blind) ANOVA, main effect of lexicality: *F* _(2, 72)_ = 38.6, *p* < 0.001; main effect of group: *F* _(1, 36)_ = 0.453, *p* = 0.505; lexicality by group interaction: *F* _(2, 72)_ = 0.546, *p* = 0.581) (see SI for details). In sum, the PPC and adjacent parieto-occipital cortices showed differential recruitment for Braille reading in people born blind as compared to reading of visual print in sighted people.

Lateralization of Braille is predicted by spoken language lateralization late in cortical processing hierarchy (i.e., LIFC) and by Braille-reading hand early in cortical processing hierarchy (i.e., S1)

Lateralization index (LI) analysis revealed left-lateralized responses to written and spoken words in vOTC and IFC of sighted readers, consistent with prior studies (one-sample *t* tests of LI = 0, visual print words > rest: vOTC: *t*_(18)_ = 5.235, *p* < 0.001; IFC: *t*_(17)_ = 3.854, *p* < 0.001; spoken words > rest: vOTC: *t*_(18)_ = 2.868, *p* < 0.01; IFC: *t*_(17)_ = 2.06, *p* = 0.055; see SI for other ROIs). By contrast, the blind group did not show systematic left-lateralization in any region (one-sample *t* tests of LI = 0, Braille words > rest: vOTC: *t*_(18)_ = 0.935, *p* = 0.362; IFC: *t*_(18)_ = 0.19, *p* = 0.852; spoken words > rest: vOTC: *t*_(18)_ = 0.583, *p* = 0.567; IFC: *t*_(18)_ = -0.834, *p* = 0.415; see SI for other regions). Consistent with prior evidence, there was substantial variability in lateralization of spoken and written language among blind participants, with some participants showing strong left and others strong right lateralization (Lane et al. 2017) (Figure 3). Individual difference analysis revealed a strong relationship between lateralization of spoken and written language in the blind group in all regions except early somatosensory cortex (SMC), including IFC, vOTC and PPC (multiple regression with the LI of spoken words in IFC and dominant reading hand entered as the regressors in each region, LI of spoken words, IFC: *t*_(18)_ = 6.551, *p* < 0.001; vOTC: *t*_(18)_ = 5.23, *p* < 0.001; PPC: IFC: *t*_(18)_ = 5.83, *p* < 0.001; SMC: *t*_(18)_ = 1.684, *p* = 0.112). Conversely, relative to other regions, the SMC showed a strong effect of reading hand on lateralization (multiple regression, for the regressor dominant reading hand: *t*_(18)_ = 3.034, *p* < 0.01). The effect of reading hand was also observed in PPC and vOTC but not in the IFC (multiple regression, effect of reading hand: PPC *t*_(18)_ = 3.814, *p* < 0.01; vOTC *t*_(18)_ = 3.424, *p* < 0.01; IFC *t*_(18)_ = -0.693, *p* = 0.498, see SI for details.) Correlations between spoken and written language lateralization were weaker in the sighted group and only reached significance in IFC and PPC (IFC: *r* = 0.865, *p* < 0.001; PPC: *r* = 0.60, *p* < 0.05). This is likely due to low variability of laterality scores in the current sighted sample (i.e., uniformly strong left-lateralization, see SI for details).

**Figure 3.**
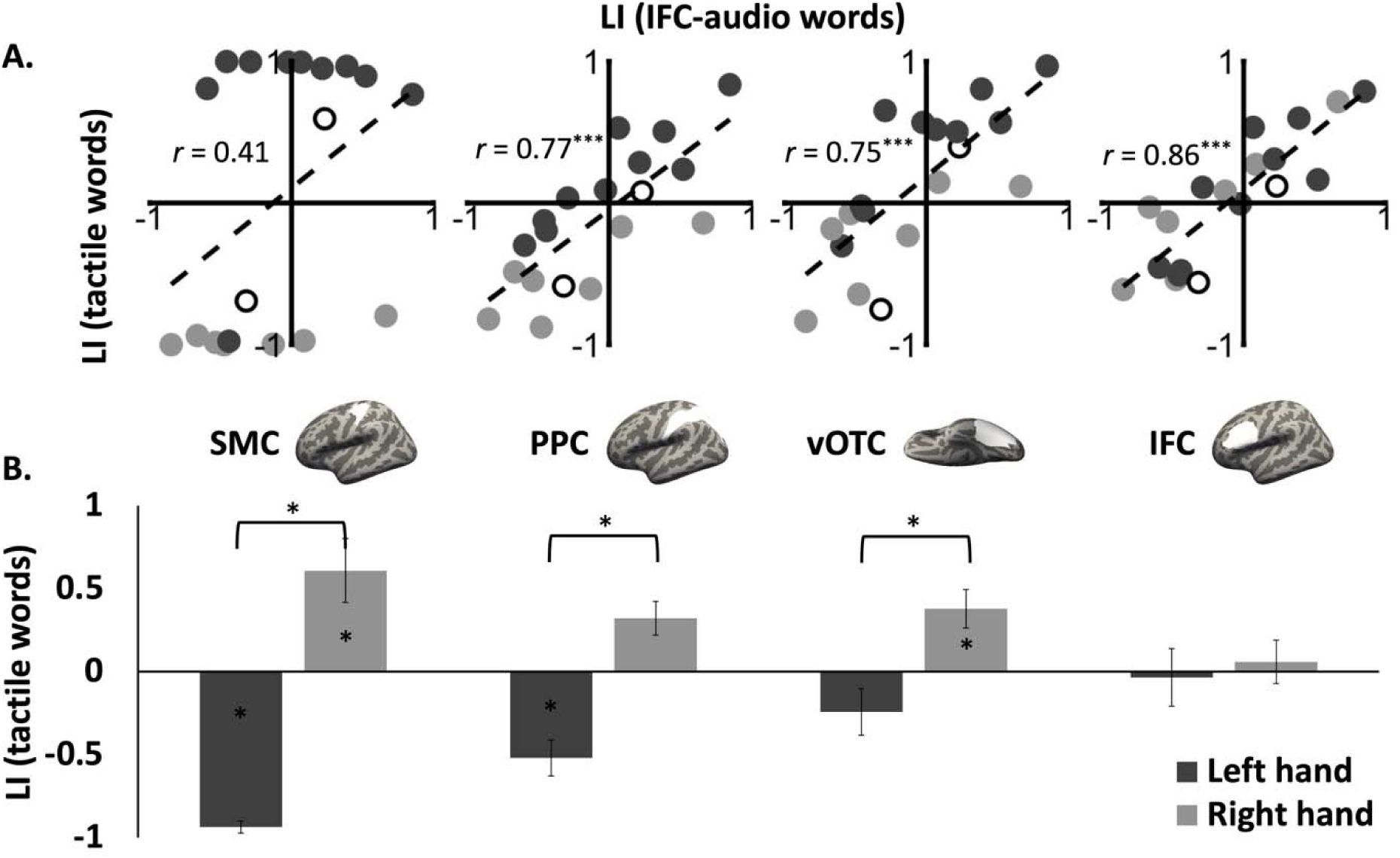
(A) Correlations between the Laterality Index (LI) of audio words in IFC and the LI of tactile words in SMC, PPC, vOTC in blind Braille readers. Data points represent individual participants, left-handed Braille readers: dark grey circles; right-handed readers: light gray circles; two subjects who did not obtain reading hand information: white circles. LI 1 score indicates strong left lateralization and -1 indicates strong right lateralization. (B) LI of Braille reading with left (grey) and right (white) hand separately. Asterisks (*) on the bar denotes significant difference from 0; asterisks (*) between two bars denote significant difference between the LI of left- vs. right-hand reading (*p* <0.05).

#### Whole cortex analyses

Reading-related activity (relative to rest) was left-lateralized in the sighted and bilateral in the blind group. For reading as compared to rest, both sighted (visual words) and blind (Braille words) readers activated the bilateral vOTC (blind peak: -40, -57, -13; sighted peak: -35, -45, -20), including the location of the classic VWFA (peak: -46, -53, -20), as well as early visual cortices, specifically the foveal confluence (V1/V2/V3) (Figure 4). vOTC responses in the blind group extended medially and anteriorly, as well as into lateral occipito-temporal cortex. Both groups also activated prefrontal cortices (inferior frontal gyrus and middle frontal gyrus). (See Table S1 for complete list of foci.)

**Figure 4.**
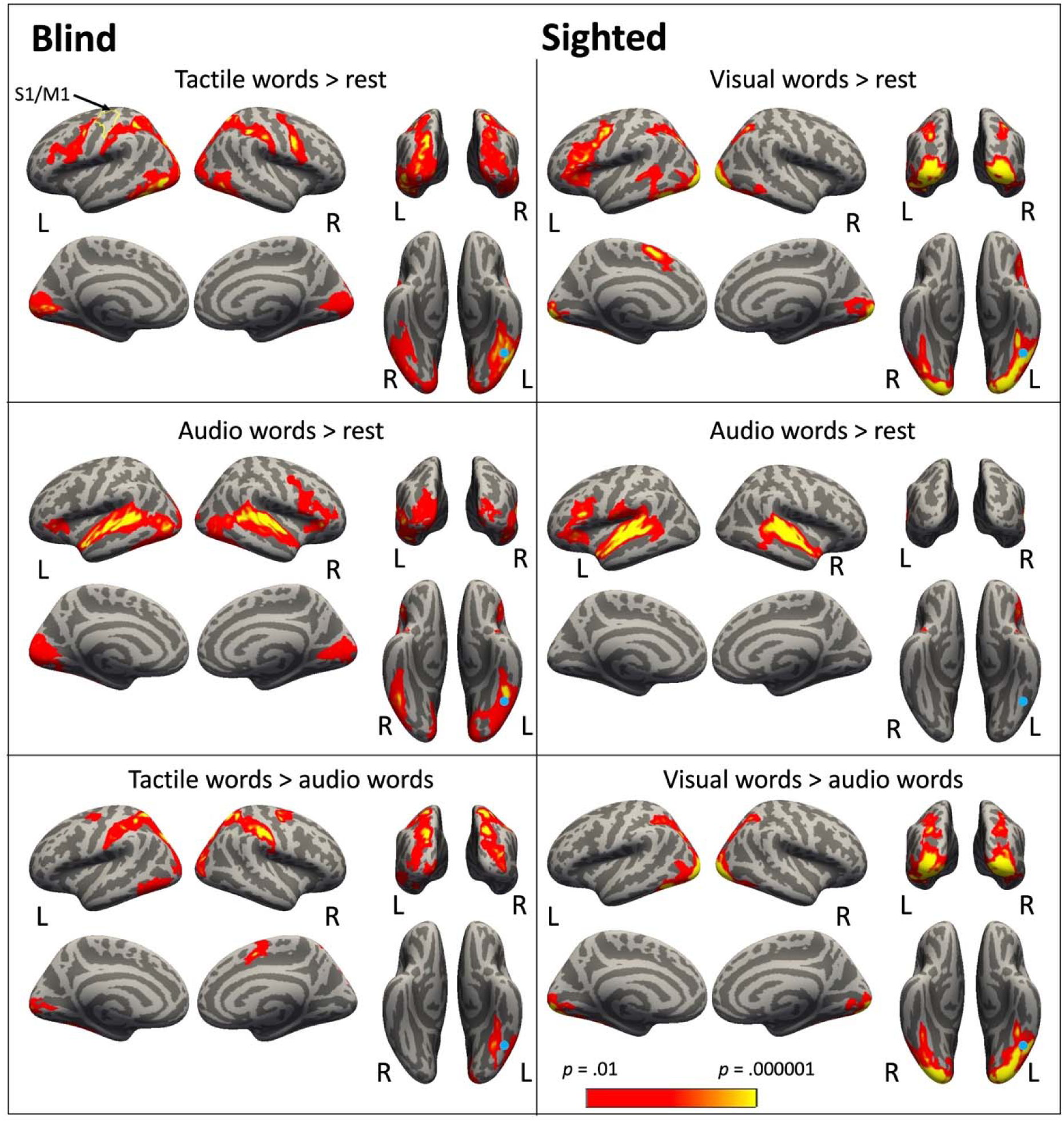
Whole-cortex results for blind (left column) and sighted (right column) *p* < 0.05 cluster-corrected. Blue circles mark previously reported location of VWFA (MNI coordinate: -46, -53, -20) (McCandliss et al. 2003). The yellow outline marks the hand S1/M1 region.

Reading Braille in the blind group (relative to rest) extensively activated posterior parietal cortices (superior parietal lobule, supramarginal gyrus (SMG)), posterior to early sensory-motor hand representations. This activity extended into parieto-occipital and dorsal occipital regions in the blind group. The sighted group activated a small cluster within PPC. A lateral temporal region was active in the sighted but not blind group.

Like responses to Braille, responses to spoken words were left-lateralized in the sighted group and bilateral in the blind group. Listening to words (audio words > rest) activated bilateral vOTC (peak: - 42, -43, -16), including the location of the classic VWFA, and early ‘visual’ cortices, in the blind group only. Both groups activated classic fronto-temporal language regions in inferior and prefrontal as well as lateral temporal cortices (Figure 4).

When reading and listening to words was compared directly, for the sighted group, reading induced greater activation in bilateral vOTC, including the typical location of the VWFA and regions posterior to it, as well as bilateral early visual cortices. The blind group also activated a region in left vOTC (fusiform; peak: -31, -58, -16), but this activation was medial to the typical VWFA location. A cluster of activity was also observed posterior to the typical VWFA location in the blind group, in the inferior temporal/lateral occipital cortex (peak: -42 -74 -5) as well as in left foveal early ‘visual’ cortices. The blind group showed more extensive activation than sighted group for Braille words > spoken words in PPC, including the SMG and superior parietal lobule, extending into dorsal occipital/parieto-occipital cortices.

In sum, although both groups activated vOTC during reading, the peak location, distribution and functional profile of responses in vOTC were distinct across groups. The blind group activated more extensive posterior parietal, parieto-occipital, and dorsal occipital areas during reading (Braille) (see Figure S8 for other contrast and Figure S9 for group comparison for all the contrast).

For the words compared to control shapes, the sighted group’s whole brain results showed stronger responses in the expected left-lateralized lateral temporal language areas, as well as left sensory-motor cortex, left medial superior frontal gyrus and bilateral early visual cortices and precuneus (see Figure S8b). The blind group’s whole brain results also showed stronger response to words over control shapes in lateral temporal language areas, extending on the lateral surface into visual cortices, as well as inferior frontal cortices. In the blind group, words>control also elicited stronger activation in bilateral vOTC and left early visual cortices. Words also elicited larger responses than consonant strings in lateral temporal cortices in both groups. In the blind readers, these responses were more posterior (see Supplementary Figure S8b for details). These whole-brain results demonstrate that although in the sighted group only small portions of vOTC responded and no portion of PPC preferentially responded to words, this is not because such effects were precluded by the current stimuli or task. We observed robust preferential responses to words in large swaths of expected, left-lateralized temporal lobe language areas.

## Discussion

### No word-form gradient in vOTC of blind Braille readers

Consistent with past research, we observed a posterior-to-anterior functional gradient, progressively higher responses to words anteriorly in the left vOTC of sighted readers. The posterior-most portion of vOTC showed no preference for words, consonant string or false font, the middle for consonants and the anterior-most showed an equivalently high response to words and consonants (Bruno et al. 2008; Cohen et al. 2008; Dehaene and Cohen 2011; Ludersdorfer et al. 2013). By contrast, in the vOTC of blind readers we found no evidence for left-lateralization, no evidence for a posterior-to-anterior functional gradient, or posterior/anterior change in modality preference. The entire posterior/anterior extent of bilateral vOTC, as well as V1, showed a preference for words in the blind group (Sadato et al. 1996; Cohen et al. 1997; Kupers et al. 2007). There was also no change in preference for written as opposed to spoken words along the posterior/anterior extent of vOTC in blind readers (see Figure S2 in SI). In sum, we find no evidence for an orthographic gradient in vOTC of blind Braille readers.

Although we found no word-form gradient in vOTC of blind Braille readers, the canonical location of the VWFA responded to words in both blind and sighted groups. Even the functional profile of the VWFA location differed somewhat across groups, however. In sighted readers the canonical location of the VWFA responded as much or more to consonant strings as to words in the current task. This is consistent with what was reported in a previous study with a smaller subsample of the present participants (Kim et al. 2017). Importantly, this pattern of VWFA response is also consistent with previous studies with sighted participants using attentionally demanding reading tasks analogous to the one used in the current study. Such tasks include longer presentation times or require attention to the orthographic content of the stimuli, like one-back memory task or phonological task (Bruno et al. 2008; Dehaene and Cohen 2011; Twomey et al. 2011; Ludersdorfer et al. 2013; Mano et al. 2013). Like these prior studies, the current experiment used relatively long presentations and involved a working memory task. Under such conditions, the VWFA responds as much or more to non-words and letter strings as to words. This response profile has been interpreted to reflect the increased demands on orthographic processing for letter strings and nonwords relative to words in tasks that require attention to orthography. By contrast, in studies using tasks that are relatively quick and automatic, like passive reading or visual feature detection, the VWFA responds more to words than consonant strings or pseudowords, putatively because more anterior portions of the VWFA code letter combinations and whole words (Cohen et al. 2002; Binder et al. 2006; Vinckier et al. 2007).

In the current study, which used an attentionally demanding task and relatively slow presentation, the VWFA of blind readers responded more to words than consonant strings, whereas that of sighted readers responded equally or more to consonant strings. One interpretation of this difference in VWFA profile across groups, is that the VWFA supports orthographic processing in the sighted but is sensitive to high level linguistic information (semantic, syntactic) in people born blind. No amount of attention to orthography can override this preference. Regardless of the interpretation of this particular group difference, we find that the same conditions that are sufficient to elicit an orthographic gradient response in the sighted do not do so in people who are blind.

Whole-cortex analysis also revealed a partially distinct distribution of reading responses in vOTC across blind and sighted people. When Braille and spoken words were each compared to rest, a peak of activation was observed in the VWFA location in both groups. However, when reading words was compared to hearing words, activity peaked medial to the VWFA in the blind but not sighted group (peak: -27, -61, -14). A similar medial vOTC region was recently found to be functionally connected with dorsal parietal cortices in sighted people (Bouhali et al., 2019; Cohen et al., 2008; Corbetta & Shulman, 2002; Henry et al., 2005; Saalmann et al., 2007). Previous studies have also found that in people who are blind, the classic lateral VWFA location is sensitive to syntactic complexity of spoken sentences, shows enhanced responses to spoken language and enhanced connectivity with fronto-temporal language networks (Burton, Snyder, Diamond, et al. 2002; Striem-Amit et al. 2012; Lane et al. 2015; Kim et al. 2017; Dzięgiel-Fivet et al. 2021). An intriguing possibility to be tested in future work is that medial vOTC is responsive to Braille-specific input from PPC, whereas the classic, more lateral VWFA location, is driven by linguistic information from fronto-temporal networks.

### Parieto-occipital decoding stream in blind readers of Braille

We observed more extensive and different involvement of parietal and parieto-occipital regions in Braille as opposed to visual print reading. Some past studies have also reported parietal activity during Braille reading but had not explored whether these responses were Braille-specific or merely related to general tactile perception (Sadato et al. 1998; Burton, Snyder, Conturo, et al. 2002; Burton et al. 2012; Siuda-Krzywicka et al. 2016; Dzięgiel-Fivet et al. 2021). Using sensitive individual-subject analyses, we found that in blind readers, subregions of posterior PPC and parieto-occipital cortices respond preferentially to Braille words relative to both tactile shapes and spoken words. This functional profile is consistent with its specific involvement in Braille reading. In contrast, anterior regions of PPC, adjacent to S1, responded most to unfamiliar tactile shapes comprised of Braille dots. The hand regions of S1 itself did not show robust or preferential responses to Braille (Burton, Snyder, Conturo, et al. 2002; Kupers et al. 2007). This response profile is suggestive of an anterior-to-posterior Braille reading processing stream, with anterior regions supporting recognition of tactile patterns and posterior regions performing Braille-specific, orthographic processing.

Further work is needed to uncover the precise contribution of PPC and parieto-occipital cortices to Braille reading. In sighted readers, the PPC shows sensitivity to phonological rather than orthographic information, during visual reading, and is involved in effortful letter-by-letter reading (e.g., when words are degraded) (Booth et al. 2003; Henry et al. 2005; Costanzo et al. 2012; Taylor et al. 2013; Ossmy et al. 2014; Bouhali et al. 2019) . Whether the same portions of PPC contribute to Braille and visual reading is not known and the precise contribution of PPC to Braille and visual reading remains to be fully characterized. It is worth noting that the Braille words used in the current study were written in Grade-II contracted Braille, which is the most used form of Braille in the US and therefore the ecologically valid choice. About 60% of the Braille word stimuli in the current study contained one or more contractions, i.e., a single Braille character that stands for multiple letters. We would argue that a full understanding of the neural basis of Braille reading requires investigating Braille reading as it is naturally done by proficient readers. How Braille contractions influence neural responses in the reading network is not known and should be examined in future research. It is possible that some of the differences between the neural basis of Braille and the neural basis of visual print relate to the contracted nature of Braille. For example, it would be worth testing the role of PPC in unpacking Braille contractions. Given previously documented similarities between the neural bases of various visual scripts, we think it is unlikely that most of the robust group differences we observed in the current study are related to the contracted nature of Braille alone. Nevertheless, this topic merits further investigation.

### Lateralization of Braille reading: effects of spoken language laterality and reading hand

We observed reduced and variable lateralization of both written and spoken language networks in blind compared to the sighted individuals. This observation is consistent with prior studies using spoken sentences (Lane et al., 2017; also see Röder et al., 2002). The reason for reduced lateralization of spoken language individuals born blind is not known. One hypothesis is that in sighted individuals language is “forced” out of the right hemisphere by visuospatial functions and such pressure is different in people born blind (Levy 1969; Kosslyn 1987). Alternatively, changes in the timing of language acquisition could affect their lateralization patterns in blindness (Locke 1997; Bishop 2013).

Importantly, here we find that laterality of responses to written words are predicted by the laterality of spoken language across blind individuals and this effect increases along the processing hierarchy peaking in IFC. In other words, those blind individuals who show right-lateralized responses to spoken words also show right-lateralized responses to written words. The only region which did not show a relationship between laterality of spoken and written language is early somatosensory cortex, where laterality was predicted only by reading hand. Previous studies with sighted readers with right hemisphere spoken language responses have likewise observed co-lateralization of spoken and written language, although this relationship was weak in the current study, likely due to low laterality variability in our sighted sample (Cai et al. 2010; Van der Haegen et al. 2012). The striking relationship between lateralization of Braille and spoken language suggests that written and spoken language co-lateralize regardless of reading modality. This observation is consistent with the hypothesis that strong connectivity and proximity to spoken language networks is one of the determining factors of which regions become ‘recycled’ for reading (Hannagan et al. 2015; Saygin et al. 2016; Stevens et al. 2017).

In contrast to the effect of spoken language on laterality, the effect of reading hand was strongest at early stages of processing (in the primary somatosensory cortex), weaker at intermediate stages (in PPC and vOTC), and absent in a high-level language region (IFC). These laterality effects support the view that PPC and vOTC participate in reading related processes in blindness, rather than solely sensory recognition or high-level language processing.

It is worth noting that in the current study, the participants were asked to read single words with one hand. However, in naturalistic reading, where multiple lines of text need to be scanned, readers often use both hands (Millar 2003). The precise contribution of each hands is not entirely clear (Hermelin and O’connor 1971; Millar 2003). Typically one hand is the dominant reading hand whereas the other is used to run ahead to track position along the page and gain a preview (Millar 1984, 2003). The neural basis of naturalistic two-handed reading remains to be investigated in future research.

## General conclusions

The neural basis of tactile Braille reading in congenitally blind individuals and visual print reading in sighted people is governed by analogous connectivity principles but has distinct anatomical profiles. While visual print reading recruits a posterior/anterior vOTC gradient, no such gradient is observed in the vOTC of blind readers of Braille. Blind readers of Braille recruit posterior parietal cortices to a greater degree and in a different way compared to visual print reading in sighted people. Only blind readers show preferential responses to written words in PPC and parieto-occipital cortex. We observed suggestive evidence for an anterior-to-posterior stream of processing in the parietal cortex of blind Braille readers, with anterior parietal areas responsive to non-Braille tactile patterns and more posterior parietal, parieto- occipital and dorsal occipital regions responsive to Braille words. In blind and sighted readers alike, lateralization of spoken language predicts lateralization of written language. In blind readers of Braille, the effect of spoken language on laterality of Braille becomes more pronounced along the cortical processing hierarchy. Conversely, reading-hand affects lateralization more in lower stages of processing. Together these results suggest that analogous connectivity principles govern neural instantiation of reading in sighted and blind readers: the localization of reading depends jointly on connectivity to sensory input regions (unilateral S1/ bilateral V1) and language networks. These principles, however, give rise to distinct anatomical profiles depending on modality of reading and neural basis of spoken language in a particular individual.

## Data and Code Availability Statement

Experimental stimuli and code are available on https://osf.io/u2akn/. fMRI analyses utilize tools from FSL version 5.0.9 (https://fsl.fmrib.ox.ac.uk/fsldownloads/oldversions/), Freesurfer version 5.3.0 (https://surfer.nmr.mgh.harvard.edu/pub/dist/freesurfer/5.3.0/), Connectome Workbench version 1.2.0 (https://github.com/Washington-University/workbench/releases/tag/v1.2.0), and Python 2.7 (https://www.python.org/download/releases/2.7/) as well as in-house fMRI analysis software (https://github.com/NPDL/NPDL-scripts)

The whole-brain group maps of results are available at https://neurovault.org/collections/10923/. IRB protocols do not currently permit sharing of fMRI data from individual participants. We are working on the IRB permission to publicly post de identified data. If and when permission is granted by IRB, we will post the de-identified data from this study to https://openneuro.org/. Individuals seeking access to raw data should contact Dr. Marina Bedny.

## Credit Author Statement

J.K., S.K., and M.B. designed research; J.K., S.K., and M.B. performed research; M.T. analyzed data; M.T., E.S. and M.B. wrote the paper.

## Acknowledgments

We would like to thank all our blind and sighted participants, the blind community and the National Federation of the Blind. Without their support, this study would not be possible. We would also like to thank the F. M. Kirby Research Center for Functional Brain Imaging at the Kennedy Krieger Institute for their assistance in data collection.

