## Supplemental material for "Sensory Modality and Spoken Language Shape Reading Network in Blind Readers of Braille"

**This PDF file includes:**

Supplementary Method

Supplementary Results

Figures S1 to S10

Table S1

SI References

**Supplementary Method**

### Stimuli

The word stimuli consisted of 240 common nouns, verbs, and adjectives. Each participant was presented with 120 of the 240 words during the reading task; the other 120 words were presented auditorily during the listening task. The word lists were counterbalanced across participants. The Braille characters contain between 1-6 raised pins in set positions within a 2 x 3 array. In Grade-II contracted English Braille, there are contractions such that single Braille characters represent frequent letter combinations (e.g., “th”) or frequent whole words (e.g., the “c” can stand for “can”). With contractions, the Braille words were on average 4 Braille characters (range = 1-8 Braille characters, SD = 2.1 characters) and 11 tactile pins per word. In the tactile consonant string condition, there were 24 strings repeated 5 times throughout the experiment. Each string stimulus consisted of 4 Braille letters, which were created using 20 English consonants. Last, the tactile control stimuli consisted of 24 unique strings of 4 non-letter shapes made of Braille pins. Note that any dot array within a 2 x 3 grid could be part of a Braille character. Therefore, to prevent participants from processing the shapes as Braille letters, the shapes varied in size and pin number within arrays ranging in size from 4 × 5 to 7 × 7. The average number of Braille pins per string in the control condition was 58.

For the sighted group, the word stimuli consisted of 240 common nouns, verbs, and adjectives that were on average 4 letters long (range = 3-5 letters, SD = 0.7 letters). Visual word stimuli consisted of a new set of words matched to the Braille words on average character length (i.e., 4 visual letters matched to 4 Braille characters), raw frequency per million, averaged frequency per million of orthographic neighbors, and averaged bigram frequency (all comparisons *p* > 0.4, obtained from the MCWord Orthographic Wordform Database) (Medler and Binder 2005). Different groups of words were used for the visual and Braille experiment to enable character length matching since Braille contractions represent two or more English letters with a single Braille character. The visual consonant strings were the same 24 consonant letter combinations from the tactile consonant strings described above. Lastly, the control stimuli in the visual reading task were 24 unique strings, each comprised of 4 characters, which were false fonts. There were 20 false font characters in total, which matched the 20 English consonants on the number of strokes, presence of ascenders and descenders, and the stroke thickness. The stimuli for the listening task were taken from each group’s respective word list. For the audio word condition, stimuli were 120 words taken from the reading task described above.

### Procedure

For the blind participants, each trial began with a 0.5 s auditory cue instructing participants to “Touch’ (reading trial), or “Listen” (listening trial). Then participants felt or heard blocks of 6 target items, one at a time. For 10 of the blind participants, tactile target stimuli were presented on the Braille display for 2 s, followed by a 0.75 s inter-stimulus interval (ISI) (6-item list duration: 16.5 s) (Kim et al. 2017). For the newly added 9 blind participants, the ISI was lengthened to 1.75 s due to a coding error which caused the 6-item list duration to be prolonged to 22.5 s. Control analyses revealed no effects of ISI duration on the results and the data are henceforth combined. After the 6-item list had been presented, there was a short delay (0.2 s), followed by a beep (0.5 s). Then a probe stimulus (2 s) was then presented, and participants indicated with a key press whether or not the probe had been present in the list. Participants had 5.3 s to make a response. The participants were asked to read with their dominant hand and responded with the other hand. The listening task was analogous in format to the reading task. The audio words and backward speech were on average 0.41 s long. The timing and sequence of events were identical for the listening task (6-item list duration 16.5 s).

For sighted participants, the trial event sequence (cue, 6-item block, beep, probe, response) was analogous to above. Each trial began with an auditory cue instructing participants to “Look” (reading trial) or “Listen” (listening trial). During reading trials, 6 visual stimuli appeared centrally for 1 s each, followed by an ISI of 0.75, during which participants were asked to maintain gaze on a black central fixation cross (total block duration: 10.5 s). Note that visual reading blocks were shorter than tactile reading blocks for the blind participants because pilot testing indicated that visual reading is faster under these conditions. Listening trials also had a total stimulus block duration of 10.5 s, to be consistent with the reading trials within the sighted group.

### fMRI ROI analysis

To construct the left vOTC search space, we first combined the left fusiform, inferior temporal, and lateral occipital parcels from Freesurfer’s automated aparc parcellation and then excluded V1, V2 regions, and the vertices with y-axis greater than -30 (Desikan et al. 2006; Lerma-Usabiaga et al. 2018). To test the posterior-to-anterior function gradient, the left vOTC search space was divided to three portions: posterior (y < -64), middle (-48 > y > = -64), and anterior portion (y > = -48). The search space in the right hemisphere was created by flipping the left vOTC masks along the x-axis. The V1 search space was defined from a previously published anatomical surface-based atlas (PALS-B12) (Van Essen 2005). The left inferior frontal language (IFC) search space was defined by using a sentence vs. non-words contrast from a previously published study (Fedorenko et al. 2010). The parietal search space was defined by the orthogonal contrast of all tactile conditions (words, consonant strings, and control) > rest in whole-cortex analysis, excluding the occipital parcels from Freesurfer’s automated aparc parcellation (Desikan et al. 2006).

To avoid using the same data to define ROIs and to test hypotheses, a leave-one-run-out cross-validation procedure was used. ROIs were defined based on data from all but one run, then the percent signal change (PSC) was extracted from the left-out run. This procedure was repeated iteratively across all runs and the PSC was averaged across iterations.

### Laterality index analysis

The bootstrap/histogram method was used to ensure that LIs were not overly influenced by arbitrary activation threshold choices or outlier voxels (Wilke and Schmithorst 2006). Bootstrapped LIs were computed using 20 evenly spaced thresholds ranging from *z* = 1.28 to *z* = 4.26 (corresponding to one-sided *p* = 0.1 to *p* = 0.00001, uncorrected). For every threshold, each participant’s *z* statistic map was masked to only include the voxels exceeding the threshold within the search space. Then we sampled the suprathreshold voxels 100 times with replacement in each hemisphere at a sampling ratio k = 1.0. The LIs were then calculated using each pair of left and right hemisphere samples, yielding a histogram of 10,000 threshold-specific LIs. Next, a single LI for each threshold was calculated by averaging the values after removing the upper and lower 25% of the 10,000 threshold-specific values. Finally, the LI reported for each participant represents the average across all thresholds.

A small number of participants were excluded from the LI analysis for a particular region if they did not have suprathreshold activation in both hemispheres (listening task- SMC: 2 sighted, 5 blind participants excluded; PPC: 1 sighted; V1: 6 sighted; IFC: 1 sighted; reading task- SMC: 4 sighted; PPC: 1 sighted; IFC: 1 sighted).

### The vector-of-ROI analysis

A vector-of-ROI analysis was conducted to plot the response pattern to Braille words, consonant strings, and tactile shapes in the PPC extent (Konkle and Caramazza 2013). To partition the PPC into a series of sub-ROIs, we (1) draw a series of anchor points along the PPC mask; (2) fit a spline through the anchor points; (3) redefine the anchor points so that the spline is divided into 15 segments of equal distance; (4) draw a series of sub-ROIs with 25 mm radius centered on these anchor points, and then extract the overlap between each circle and the PPC mask; (5) remove the overlap between ROIs.

For consistency, the same procedure was performed for both the left and right PPCs. However, the left and right PPCs are defined by the group contrast of all tactile stimuli > rest, then the size and shape of these two masks are different. After removing the overlap between sub-ROIs, only 13 valid sub-ROIs were defined in the left PPC. Fifteen valid sub-ROIs were defined in the right PPC. In each sub-ROI, the mean beta value for each condition was extracted for each participant.

#### Supplementary Results

#### Visual (sighted) but not tactile Braille reading (blind) elicits a posterior-to-anterior functional gradient in left vOTC and shows left-lateralization

A four-way hemisphere (left, right) by posterior/anterior subregion (posterior, middle, anterior) by lexicality (words, consonant strings, control) by group (sighted, blind) ANOVA were conducted to examine reading responses across groups.

This ANOVA revealed significant main effects of hemisphere (left hemisphere > right hemisphere, *F* _(1, 36)_ = 7.917, *p* < 0.01), posterior/anterior subregion (posterior and middle > anterior, *F* _(2, 72)_ = 10.554, *p* < 0.001), and lexicality (words and consonant strings > controls, *F* _(2, 72)_ = 15.077, *p* < 0.001). There was no group main effect (*F* _(1, 36)_ = 2.124, *p* = 0.154). For the two-way interaction, the group by lexicality (*F* _(2, 72)_ = 10.672, *p* < 0.001), group by posterior/anterior subregion (*F* _(2, 72)_ = 12.379, *p* < 0.001), hemisphere by lexicality (*F* _(2, 72)_ = 46.708, *p* < 0.001) and the posterior/anterior subregion by lexicality (*F* _(4, 144)_ = 7.393, *p* < 0.001) interaction effects were significant. There was no hemisphere by group interaction effect (*F* _(1, 36)_ = 0.291, *p* = 0.593) or hemisphere by posterior/anterior subregion interaction effect (*F* _(2, 72)_ = 0.946, *p* = 0.393). For the three-way interaction, the hemisphere by posterior/anterior subregion by group interaction (*F* _(2, 72)_ = 4.04, *p* < 0.05), the posterior/anterior subregion by lexicality by group interaction (*F* _(4, 144)_ = 2.546, *p* < 0.05), and the hemisphere by posterior/anterior subregion by lexicality interaction (*F* _(4, 144)_ = 2.825, *p* < 0.05) were all significant. The hemisphere by lexicality by group interaction (*F* _(2, 72)_ = 1.811, *p* = 0.717) was not significant. The four-way interaction was reported in the main Results section.

For the sighted group, we found the expected three-way interaction between hemisphere (left, right), posterior/anterior subregion (posterior, middle, anterior) and lexicality (words, consonant strings, control; *F* _(4, 72)_ = 4.344, *p* < 0.01). Next, we looked at each hemisphere separately in the sighted group.

In the left vOTC, there was a two-way interaction between lexicality (words, consonant strings, control) and posterior/anterior subregion (posterior, middle, anterior; *F* _(4, 72)_ = 8.237, *p* < 0.001), reflecting the expected posterior-to-anterior functional gradient. Pairwise comparisons revealed that the posterior vOTC responded similarly to all visual stimuli (all pairwise comparisons *p* > 0.05). By contrast, in middle vOTC, consonant strings elicited higher responses than both words and control stimuli (Bonferroni-corrected paired *t*-test for words vs. consonant strings: *t*_(18)_ = --2.429, *p* = 0.078; consonant strings vs. control: *t*_(18)_ = 3.787, *p* < 0.01; words vs control: *t*_(18)_ = 0.888, *p* = 0.386). In anterior vOTC, responses to words and consonant strings were both higher than control and not different from each other (Bonferroni-corrected paired *t*-test for words vs consonant strings: *t*_(18)_ = 0.064, *p* > 0.99; consonant strings vs. control: *t*_(18)_ = 4.053, *p* < 0.01; words vs. control: *t*_(18)_ = 3.99, *p* < 0.01).

In the right vOTC of the sighted group, a two-way lexicality (words, consonant strings, control) by posterior/anterior subregion (posterior, middle, anterior) ANOVA revealed no main effect of lexicality (*F* _(2, 36)_ = 0.87, *p* = 0.429) and no lexicality by subregion interaction (*F* _(4, 72)_ = 0.735, *p* = 0.517). The main effect of posterior/anterior subregion was significant (anterior and middle < posterior; *F* _(2, 36)_ = 8.201, *p* < 0.01). To summarize, these results demonstrate that in the sighted group, there was a posterior-to-anterior functional gradient for processing word form during reading in the left but not right vOTC.

Next, we examined these effects in the blind group. We conducted a three-way hemisphere (left, right) by posterior/anterior subregion (posterior, middle, anterior) by lexicality (words, consonant strings, control) ANOVA. Unlike in the sighted, there was no significant three-way interaction (*F* _(4, 72)_ = 0.747, *p* = 0.563). Although there was no interaction, we conducted a separate ANOVA testing for a lexicality effect across the posterior/anterior subregions for each hemisphere separately in order to match the analysis of the sighted group.

In the left vOTC of the blind group, all three (posterior, middle, anterior) subregions responded most to words, followed by consonant strings followed by tactile shapes (Figure 1). There was a two-way interaction between lexicality (words, consonant strings, control) and posterior/anterior subregion (posterior, middle, anterior; *F* _(4, 72)_ = 2.83, *p* < 0.05). However, the nature of this interaction was different from that observed in the sighted group. All pairwise-comparisons between conditions were significant in all three subregions (words > consonant strings > control), except the difference between words and consonant strings did not reach significance in the anterior vOTC (Bonferroni-corrected paired *t*-test for words vs. consonant strings: posterior vOTC *t*_(18)_ = 2.459, *p* = 0.089; middle vOTC: *t*_(18)_ = 2.865, *p* < 0.05; anterior vOTC: *t*_(18)_ = 2.093, *p* = 0.152; words vs. control: posterior vOTC: *t*_(18)_ = 4.859, *p* < 0.001; middle vOTC *t*_(18)_ = 7.682, *p* < 0.001; anterior vOTC: *t*_(18)_ = 5.561, *p* < 0.001; consonant strings vs. control: posterior vOTC: *t*_(18)_ = 3.102, *p* < 0.01; middle vOTC *t*_(18)_ = 4.461, *p* < 0.01; anterior vOTC: *t*_(18)_ = 5.067, *p* < 0.001).

Unlike in the sighted group, in the right hemisphere of the blind group, lexicality effects were similar to the left hemisphere. All three (posterior, middle, anterior) subregions responded most to words, followed by consonant strings followed by tactile shapes. There was also a two-way interaction between lexicality (words, consonant strings, control) and subregion (posterior, middle, anterior; *F* _(4, 72)_ = 6.019, *p* < 0.001). Pairwise comparisons showed that the posterior right vOTC responded more to words than control (*t*_(18)_ = 3.380, *p* < 0.01); the middle vOTC responded more to words than both consonant strings (*t*_(18)_ = 3.585, *p* < 0.01) and control (*t*_(18)_ = 4.032, *p* < 0.001); and the anterior vOTC responded most strongly to words and consonant strings than control stimuli (words vs. consonant strings: *t*_(18)_ = 2.325, *p* = 0.096; words vs. control: *t*_(18)_ = 5.09, *p* < 0.001; consonant strings vs. control, *t*_(18)_ = 3.902, *p* < 0.01). Other pairwise comparisons did not reach significance (posterior vOTC: words vs. consonant strings, *t*_(18)_ = 2.201, *p* = 0.123; consonant strings vs. control, *t*_(18)_ = 1.552, *p* = 0.414; middle vOTC: consonant strings vs control, *t*_(18)_ = 1.729; *p* = 0.303).

For the listening task, similar to the reading task, we conducted a four-way hemisphere (left, right) by subregion (posterior, middle, anterior) by lexicality (words, control) by group (sighted, blind) ANOVA. The four-way interaction effect with group was not significant and we, therefore, did not proceed to further analyses (*F* _(2, 72)_ = 1.357, *p* = 0.264). It is worth noting that in the sighted group, responses to auditory stimuli were below rest in posterior vOTC and above rest in the more anterior regions. This pattern was not observed in the blind group (see Figure S2).

#### V1 shows a preference for words in blind readers

As with vOTC, we first examined responses in left V1 during the reading task using the consonant strings > control functional ROIs (see Figure S5). A two-way lexicality (words, consonant strings, control) by group (sighted, blind) ANOVA revealed main effects of lexicality (*F* _(2, 72)_ = 5.883, *p* < 0.01) and group (sighted > blind, *F* _(1, 36)_ = 5.348, *p* < 0.05). There was also a significant lexicality by group interaction (*F* _(2, 72)_ = 8.034, *p* = 0.001). In the blind group, V1 responded most to words and there was no difference between consonant strings and control (Bonferroni-corrected paired *t*-test, words vs. consonant strings: *t*_(18)_ = 2.56, *p* < 0.01; words vs. control: *t*_(18)_ = 3.569, *p* < 0.01; consonant strings vs. control: *t*_(18)_ = 2.296, *p* = 0.231). In the sighted group, V1 responded more to control stimuli than consonant strings (Bonferroni-corrected paired *t*-test, *t*_(18)_ = 2.33, *p* < 0.05). There was no difference between other conditions (pairwise comparisons *p* > 0.05.) V1 responses in the blind group were similar when functional ROIs were defined using words > control (Figure S6). In the sighted group, however, a preference for words over false fonts (control) emerged in this alternative analysis (Bonferroni-corrected paired *t*-test, *t*_(18)_ = 3.176, *p* < 0.05; Figure S6). This latter result is consistent with some previous studies showing that V1/V2 responded more to words than non-letter control stimuli like scrambled words (Szwed et al. 2011, 2014).

For the listening task, the two-way lexicality (words, control) by group (sighted, blind) ANOVA showed a main effect of group (*F* _(1, 36)_ = 6.638, *p* < 0.05), with overall greater activation seen in blind than sighted V1, and a main effect of lexicality (*F* _(1, 36)_ = 4.721, *p* < 0.05). There is no interaction between the factors (*F* _(1, 36)_ = 1.259, *p* = 0.296). Notably in the sighted but not blind group, responses to both words and audio control were below rest (Figure S5). This pattern of results was the same in words > control ROI (Figure S6).

#### No preference for words or consonant strings in primary sensory-motor cortex (SMC) hand region of blind participants

We examined responses of the left SMC hand region to test whether it showed a similar preference for Braille words and consonant strings as the PPC (Figure S5). For the reading task, the two-way lexicality (words, consonant strings, control) by group (sighted, blind) ANOVA showed a main effect of lexicality (*F* _(2, 72)_ = 7.946, *p* < 0.001; functional ROIs were defined using the consonants > controls contrast), with higher responses to the consonant strings than control stimuli. Note that the responses to all stimuli were below rest in SMC in the blind group. There was no main effect of group (*F* _(1, 36)_ = 2.776, *p* = 0.104) and no group by condition interaction (*F* _(2, 72)_ = 1.922, *p* = 0.154). For the listening task, the two-way lexicality (words, control) by group (sighted, blind) ANOVA revealed a main effect of group (*F* _(1, 36)_ = 16.552, *p* < 0.001), with overall greater responses in sighted group than blind group. There was no main effect of lexicality (*F* _(1, 36)_ = 0.811, *p* = 0.374) and no interaction (*F* _(1, 36)_ = 0.001, *p* = 0.974). Results were similar when the SMC ROIs were instead defined using the words > controls contrast (Figure S6). In sum, unlike in the PPC, we found no evidence for specialization of SMC for Braille reading as compared to perception of control tactile shapes.

#### The left inferior frontal cortex (IFC) prefers word-like written and spoken stimuli across blind and sighted readers

We analyzed responses in the left IFC across groups with the prediction that this high-level language region would show similar response patterns across blind and sighted readers. Consistent with this prediction, responses were similar across groups for both tasks in the left IFC (Figure S5). For the reading task, a two-way lexicality (words, consonant strings, control) by group (sighted, blind) ANOVA revealed a significant main effect of lexicality, with larger responses for words and consonant strings over the control condition (*F* _(2, 72)_ = 38.6, *p* < 0.001; functional ROIs were defined using the words > controls contrast). Neither the main effect of group (*F* _(1, 32)_ = 0.453, *p* = 0.505) nor the interaction (*F* _(2, 72)_ = 0.546, *p* = 0.581) were significant. Likewise, for the listening task, the two-way lexicality (words, control) by group (sighted, blind) ANOVA revealed the expected main effect of lexicality (words > control; *F* _(1, 36)_ = 29.231, *p* < 0.001). There was no main effect of group (*F* _(1, 36)_ = 1.448, *p* = 0.237) and no lexicality by group interaction (*F* _(1, 36)_ = 0.05, *p* = 0.825). There was also no group by condition interaction when functional ROIs were defined using the words > controls contrast. Both groups still showed a preference for words over control stimuli and in this case, there was also a larger response to words over consonant strings in both groups (Figure S6). These results are consistent with prior studies showing similar responses to spoken and written language in the left inferior frontal cortex of blind and sighted adults.

#### Lateralization of Braille correlates with spoken language lateralization and Braille-reading hand

On average, the sighted group’s SMC and PPC activity was not systematically lateralized for written words (one-sample *t* tests of LI = 0, SMC: *t*_(14)_ = 1.061, *p* = 0.307; PPC: *t*_(17)_ = -0.335, *p* = 0.741). There was a marginal left-lateralized activation in V1 for written words (one-sample *t* tests of LI = 0, V1: *t*_(18)_ = 1.957, *p* = 0.066). For spoken words, we found right-lateralized activation in PPC and marginal right-lateralized activation in V1 for spoken words in the sighted group (one-sample *t* tests of LI = 0, PPC: *t*_(17)_ = -3.619, *p* < 0.01; V1: *t*_(12)_ = -2.159, *p* = 0.052). There were no systematic lateralization in SMC for the listening task (one-sample *t* tests of LI = 0, SMC: *t*_(17)_ = -0.691, *p* = 0.499). The blind group showed no systematic lateralization for written or spoken words in any region (one-sample *t* tests of LI = 0, reading: SMC: *t*_(18)_ = 0.148, *p* = 0.884; PPC: *t*_(18)_ = -1.096, *p* = 0.287; V1: *t*_(18)_ =1.005, *p* = 0.328; listening: SMC: *t*_(13)_ = -1.124, *p* = 0.281; PPC: *t*_(18)_ = 0.187, *p* = 0.854; V1: *t*_(18)_ = -0.001, *p* = 0.999).

We then determined if lateralization of the Braille reading network could be predicted by the laterality of spoken language and Braille reading hand across blind individuals. A multiple regression analysis was conducted in each region, with the LI of spoken words in IFC and dominant reading hand entered as the regressors and the LI of written words as the dependent variable. First, both the dominant reading hand and the LI of spoken words in IFC predicted the LI of written words in PPC, vOTC and V1 (PPC: dominant reading hand: *β* = 0.45, *p* < 0.001; LI of spoken words in IFC: *β* = 0.688, *p* < 0.001; model adjust *r*^2^ = 0.784; vOTC: dominant reading hand: *β* = 0.0.439, *p* < 0.01; LI of spoken words in IFC: *β* = 0.67, *p* = 0.001; adjust *r*^2^ = 0.714; V1: dominant reading hand: *β* = 0.362, *p* < 0.05; LI of spoken words in IFC: *β* = 0.72, *p* = 0.001; adjust *r*^2^ = 0.709). Second, in the IFC, only the LI of spoken words predicted the LI of written words (IFC: dominant reading hand: *β* = 0.089, *p* = 0.498; LI of spoken words in IFC: *β* = 0.842, *p* < 0.001; adjust *r*^2^ = 0.702). Last, we found in SMC, only the dominant reading hand predicted the LI of written words (dominant reading hand: *β* = 0.56, *p* < 0.01; LI of spoken words in IFC: *β* = 0.311, *p* = 0.112; adjust *r*^2^ = 0.406). To summarize, in blind individuals, responses to Braille written words and spoken words were co-lateralized to the same hemisphere across most of the Braille reading network, including the vOTC, V1, PPC, and the IFC. Braille reading hand also had an effect on the lateralization of Braille written words in vOTC, PPC, and SMC.

In the sighted group, the correlation between the LI of spoken words in IFC and the LI of written words in vOTC was not significant (*r* = 0.18, *p* = 0.475). In addition, there were no correlations between the LI of spoken words in IFC and the LI of written words in V1 or SMC (V1: *r* = -0.033, *p* = 0.895; SMC: *r* = 0.409, *p* = 0.13).

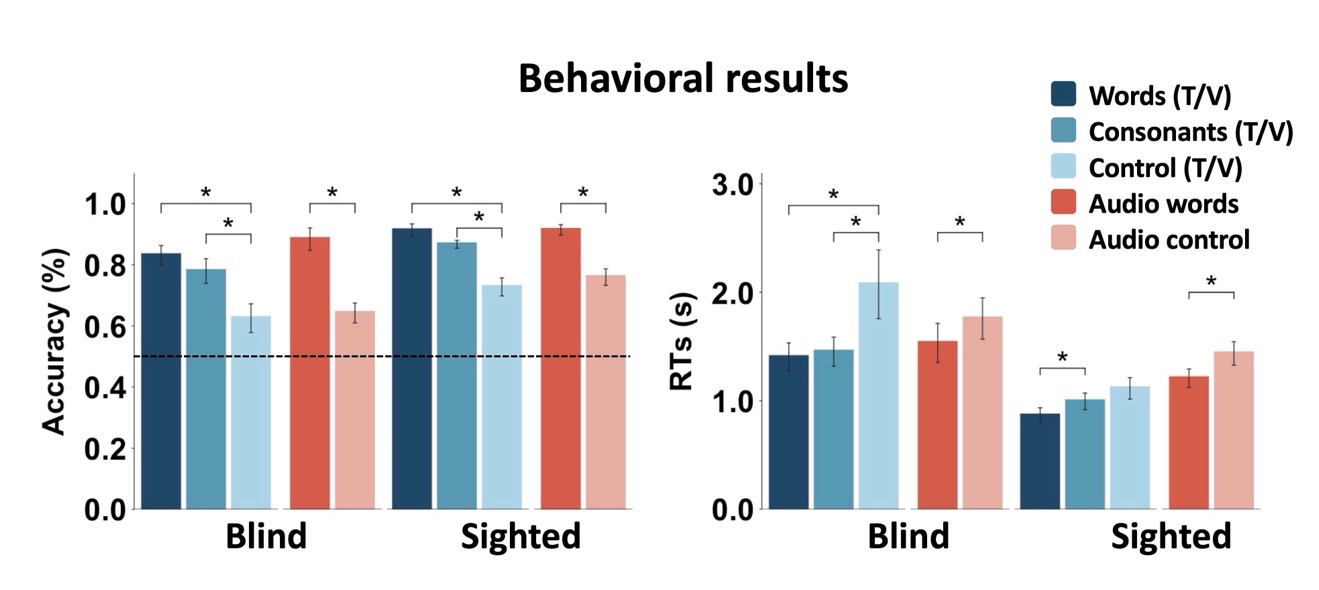

Figure. S1. Behavioral results. Accuracy (left) and reaction time (Right) results for blind (left) and sighted (right) groups for reading (blue colors) and listening (orange colors) tasks. Error bars denote standard errors +/- the mean. Asterisks (*) denote significant Bonferroni-corrected pairwise comparisons within task (*p* <.05). T = tactile, V = visual, RT = reaction time.

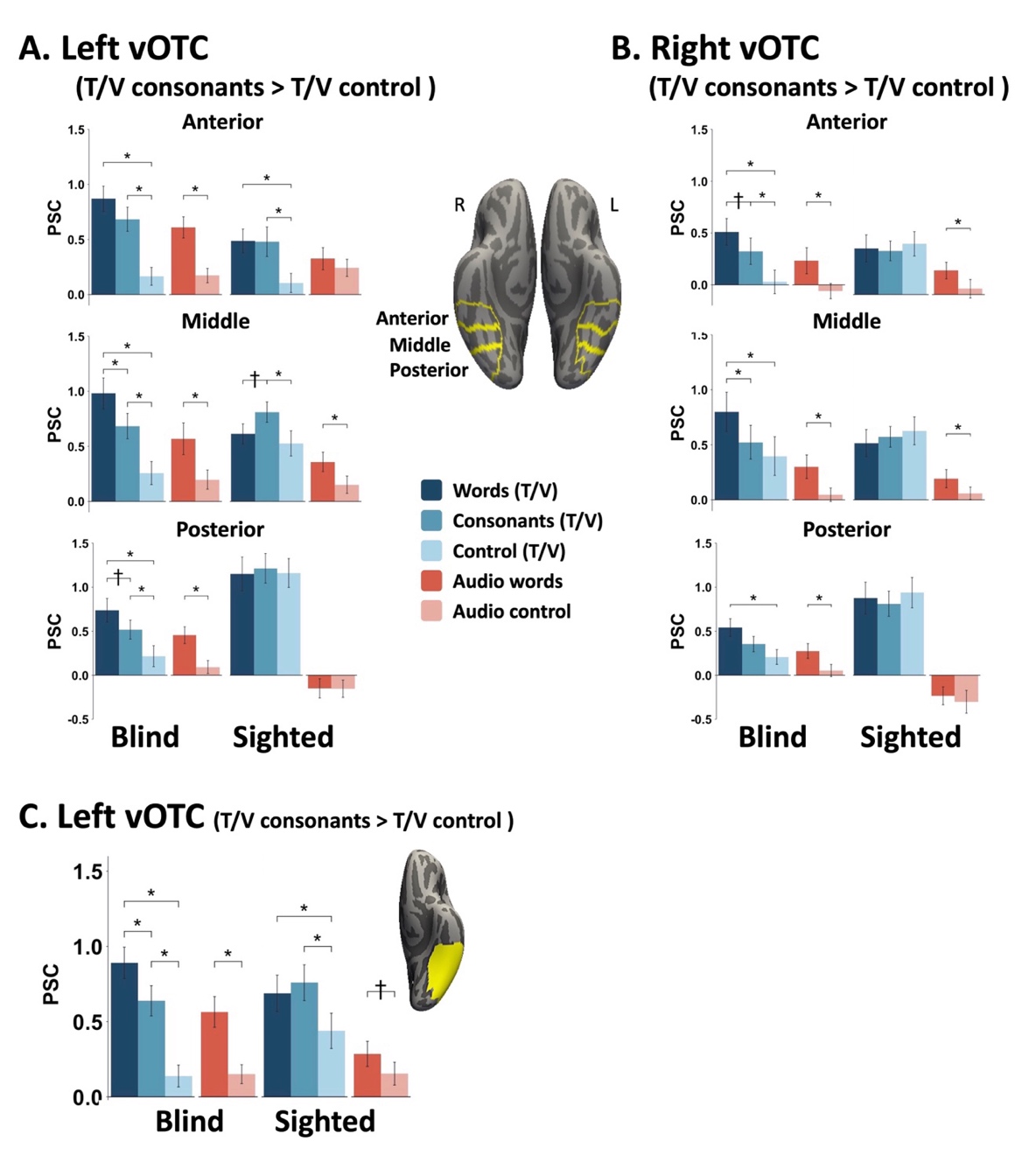

Figure. S2. Responses in (A) left and (B) right vOTC across posterior, middle and anterior subregions for blind and sighted groups during the reading (blue colors) and listening (orange colors) tasks using the consonant strings > control (false font/shape) contrast to identify individual functional ROIs (as reported in the main text). Error bars denote standard errors +/- the mean. Asterisks (*) denote significant Bonferroni-corrected pairwise comparisons within task (*p* <.05). T = tactile, V = visual.

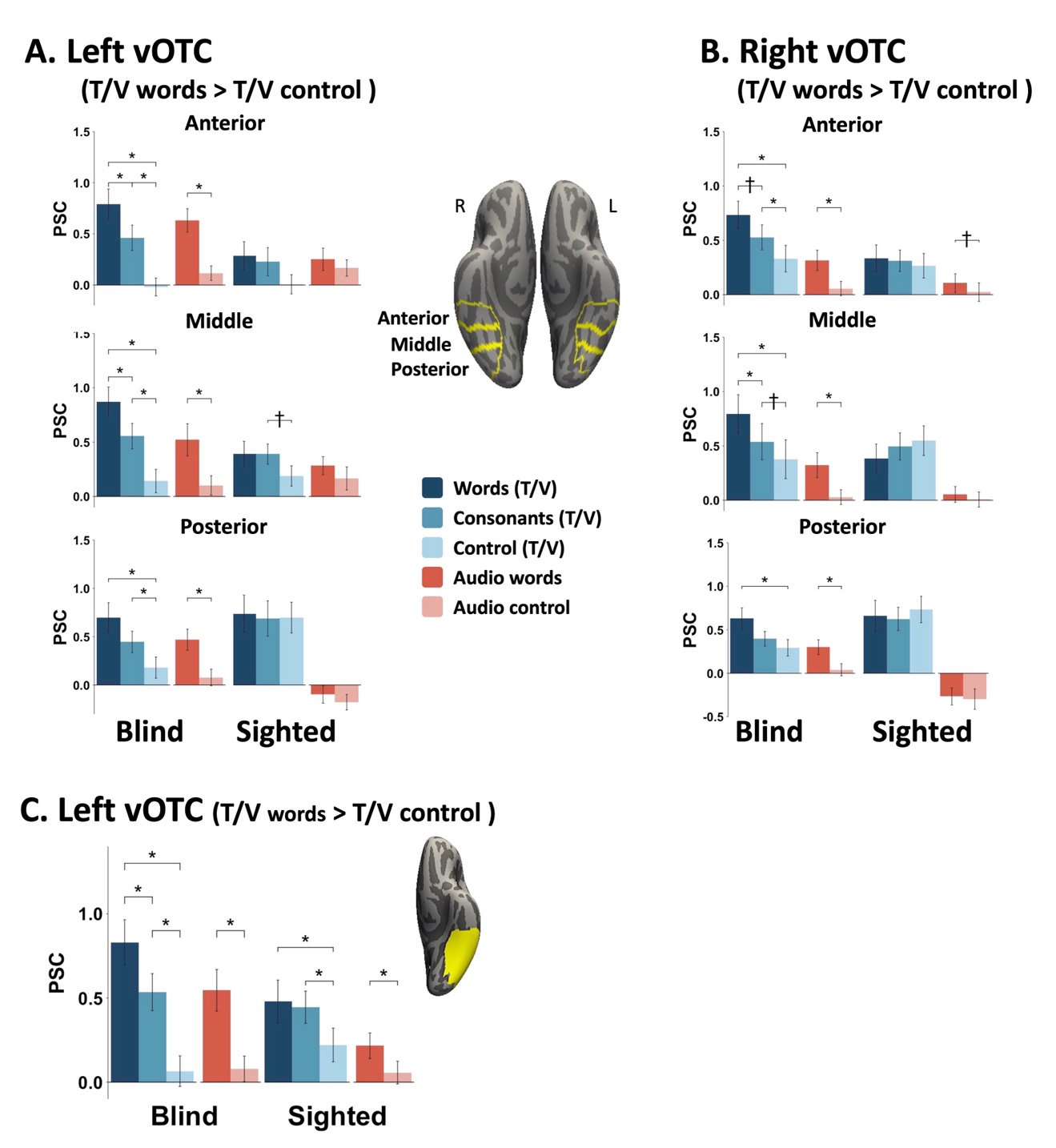

Figure. S3. Responses in (A) left and (B) right vOTC across posterior, middle and anterior subregions for blind and sighted groups during the reading (blue colors) and listening (orange colors) tasks using the words > control (false font/shape) contrast to identify individual functional ROIs. C. Error bars denote standard errors +/- the mean. Asterisks (*) denote significant Bonferroni-corrected pairwise comparisons within task (*p* <.05). T = tactile, V = visual.

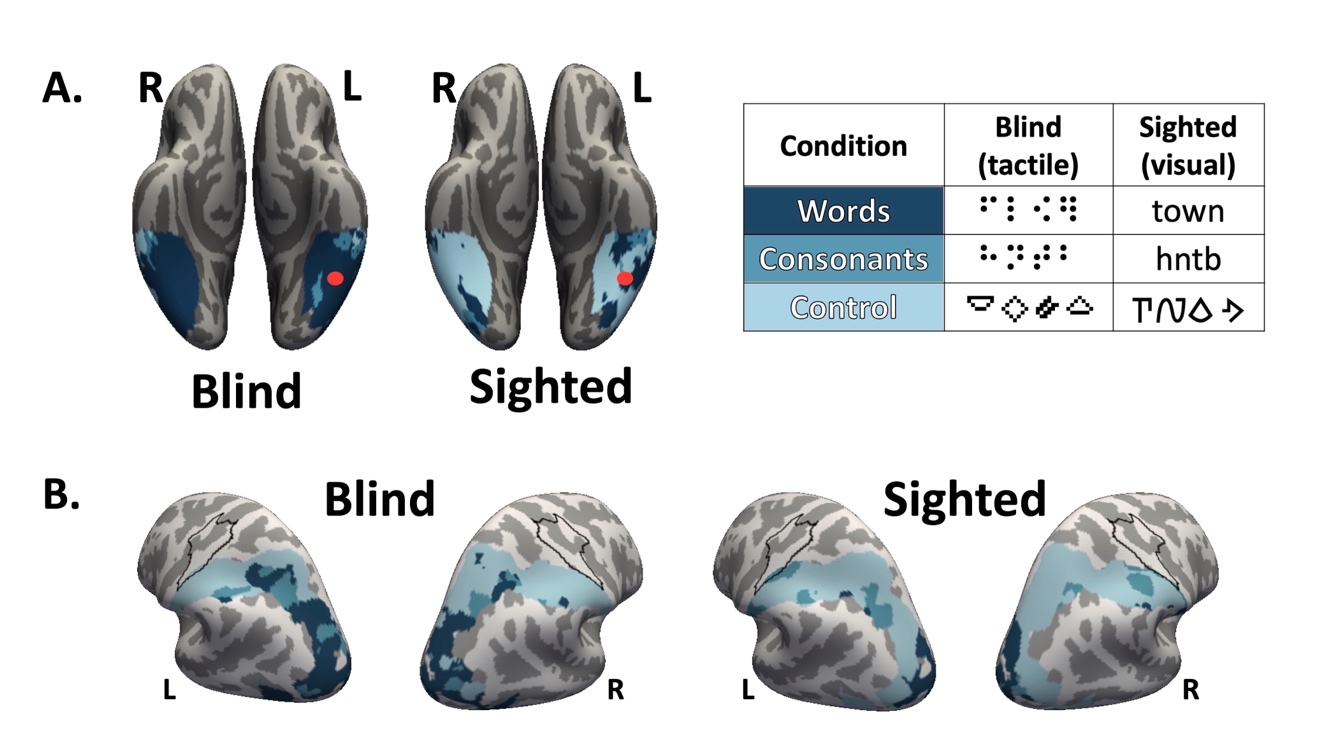

**Figure. S4.** Topographical preference winner-take-all maps of vOTC (A) and posterior parietal cortex (B) during the reading task: words, consonant strings, control stimuli. The black outline indicates the hand region of the primary sensory-motor cortex (Neurosynth). The red dots mark previously reported location of VWFA (MNI coordinate: -46, -53, -20).

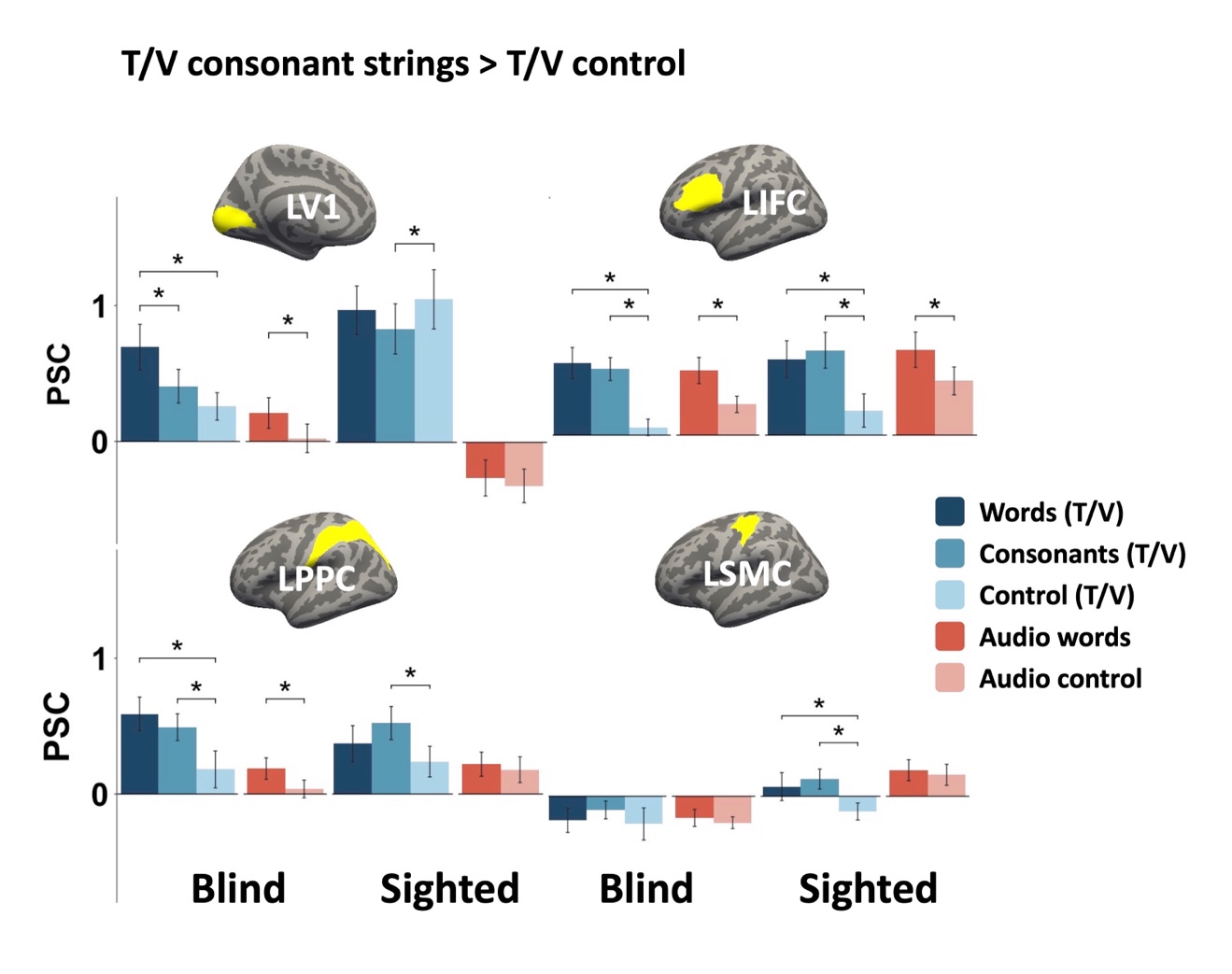

Figure. S5. Responses in left V1 (upper left), IFC (upper right), PPC (lower left), and SMC (lower right) ROIs for blind and sighted groups during the reading (blue colors) and listening (pink colors) tasks using the consonant strings > controls functional ROIs. Error bars denote standard errors +/- the mean. Asterisks (*) denote significant Bonferroni-corrected pairwise comparisons within task (p <.05). T = tactile, V = visual.

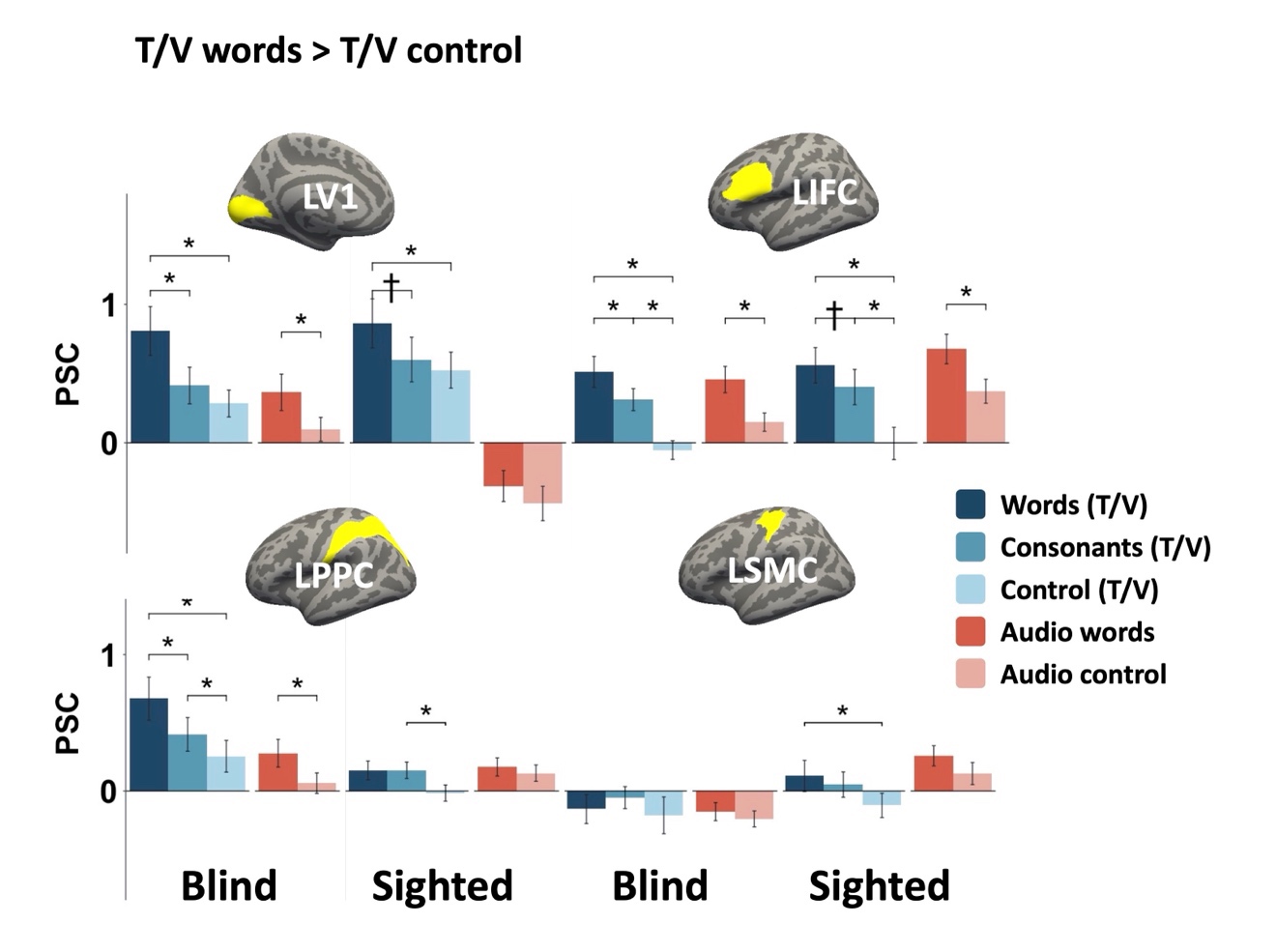

Figure. S6. Responses in left V1 (upper left), IFG (upper right), PPC (lower left), and SMC (lower right) ROIs for blind and sighted groups during the reading (blue colors) and listening (pink colors) tasks. Error bars denote standard errors +/- the mean. Asterisks (*) denote significant Bonferroni-corrected pairwise comparisons within task (p <.05). T = tactile, V = visual.

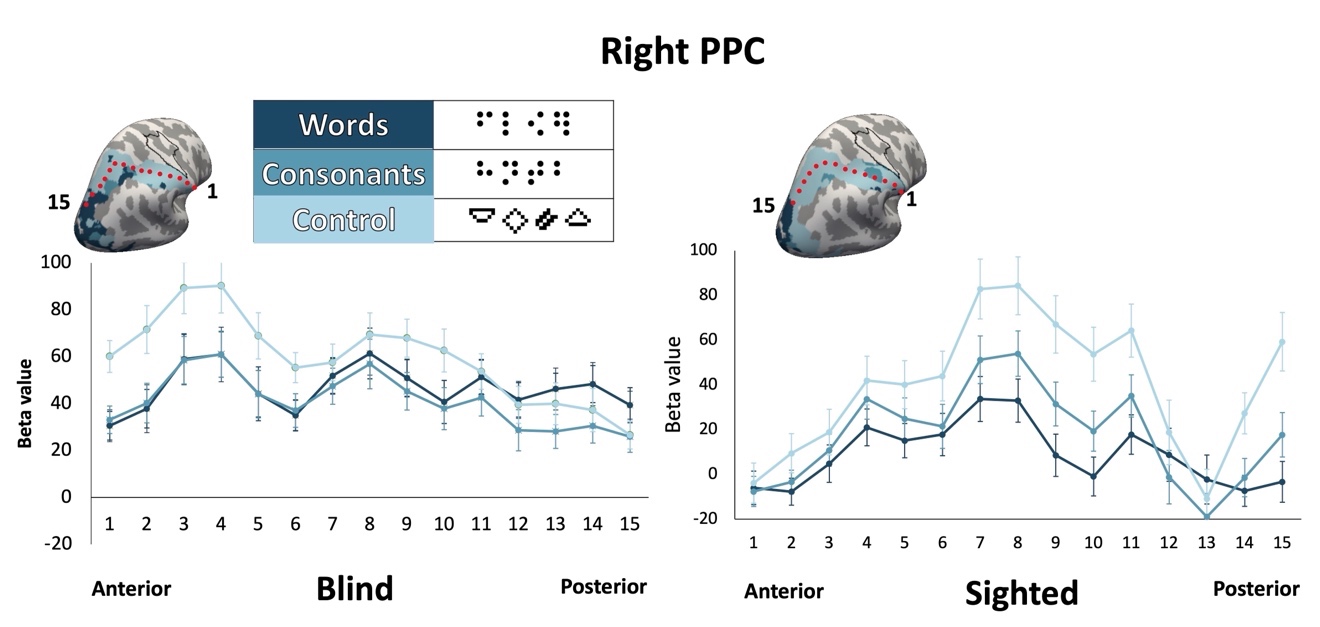

Figure. S7. Mean response to each reading condition along right anterior/posterior PPC extent in blind group (A) and left and right PPC in sighted group (B). Segment centers are marked by red dots.

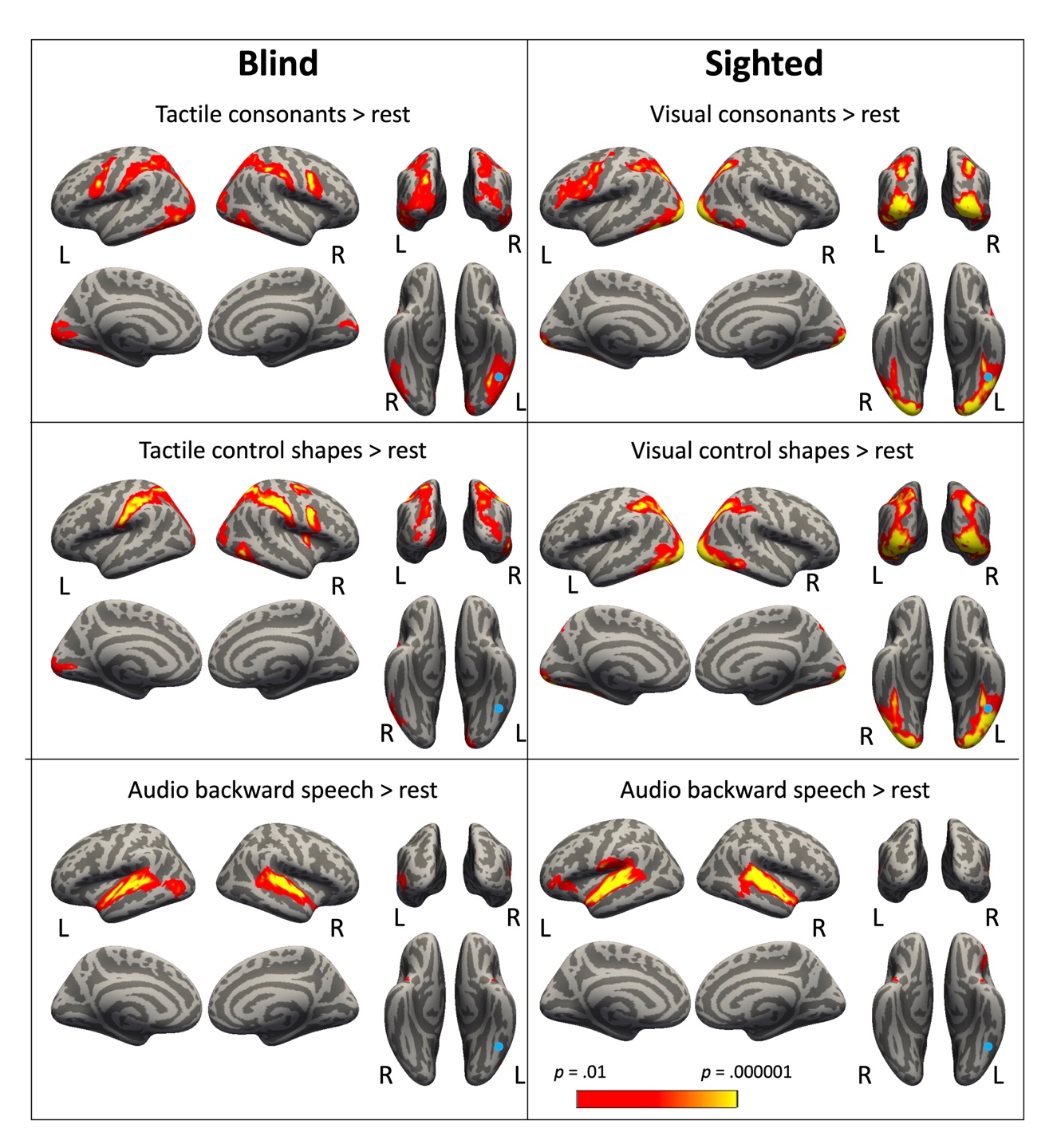

Figure. S8a. Whole-cortex results for blind (left column) and sighted (right column) *p* < 0.05 cluster-corrected. Blue circles mark previously reported location of VWFA (MNI coordinate: -46, -53, -20). The yellow outline marks the hand S1/M1 region.

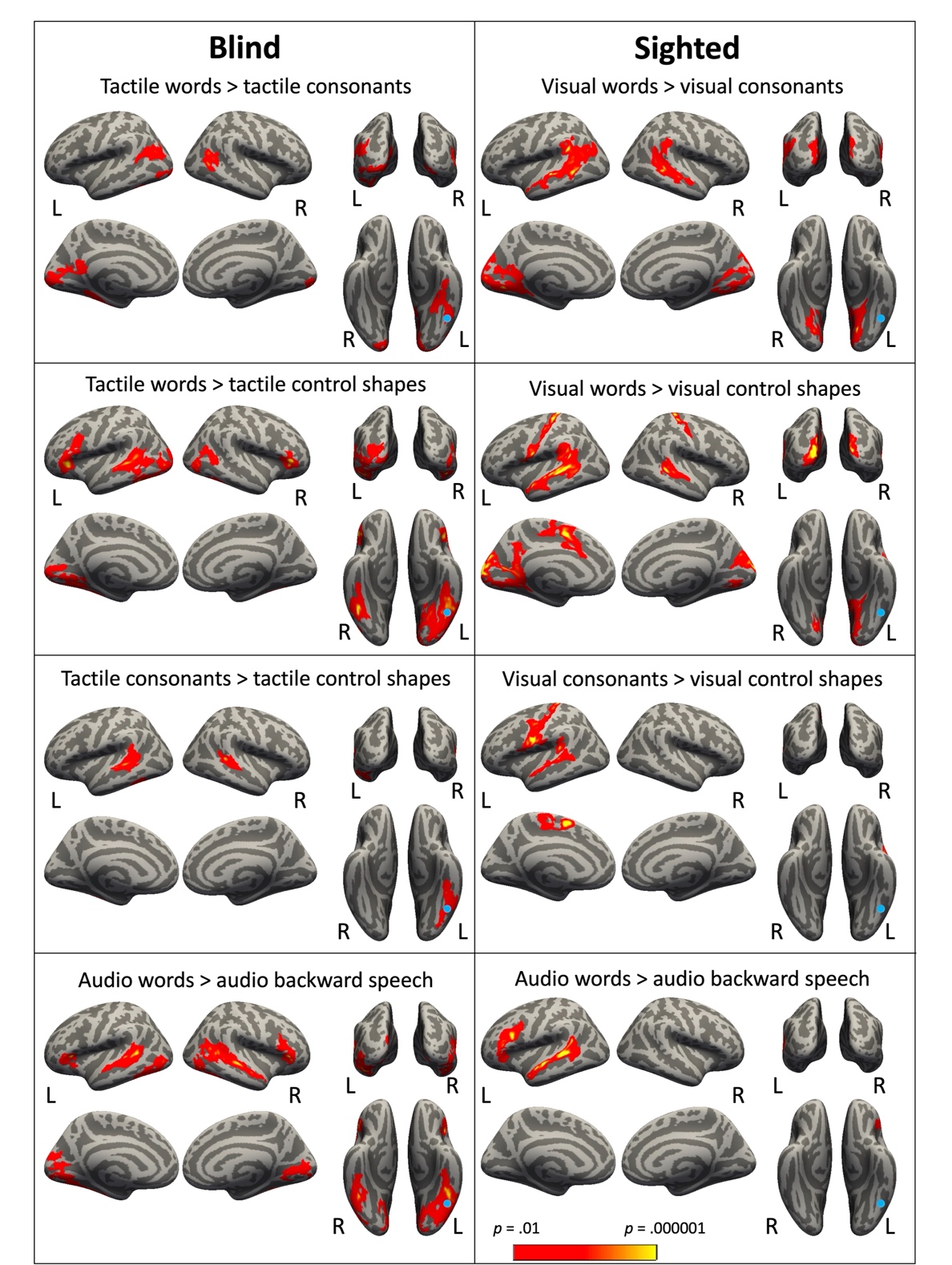

Figure. S8b. Whole-cortex results for blind (left column) and sighted (right column) *p* < 0.05 cluster-corrected. Blue circles mark previously reported location of VWFA (MNI coordinate: -46, -53, -20). The yellow outline marks the hand S1/M1 region.

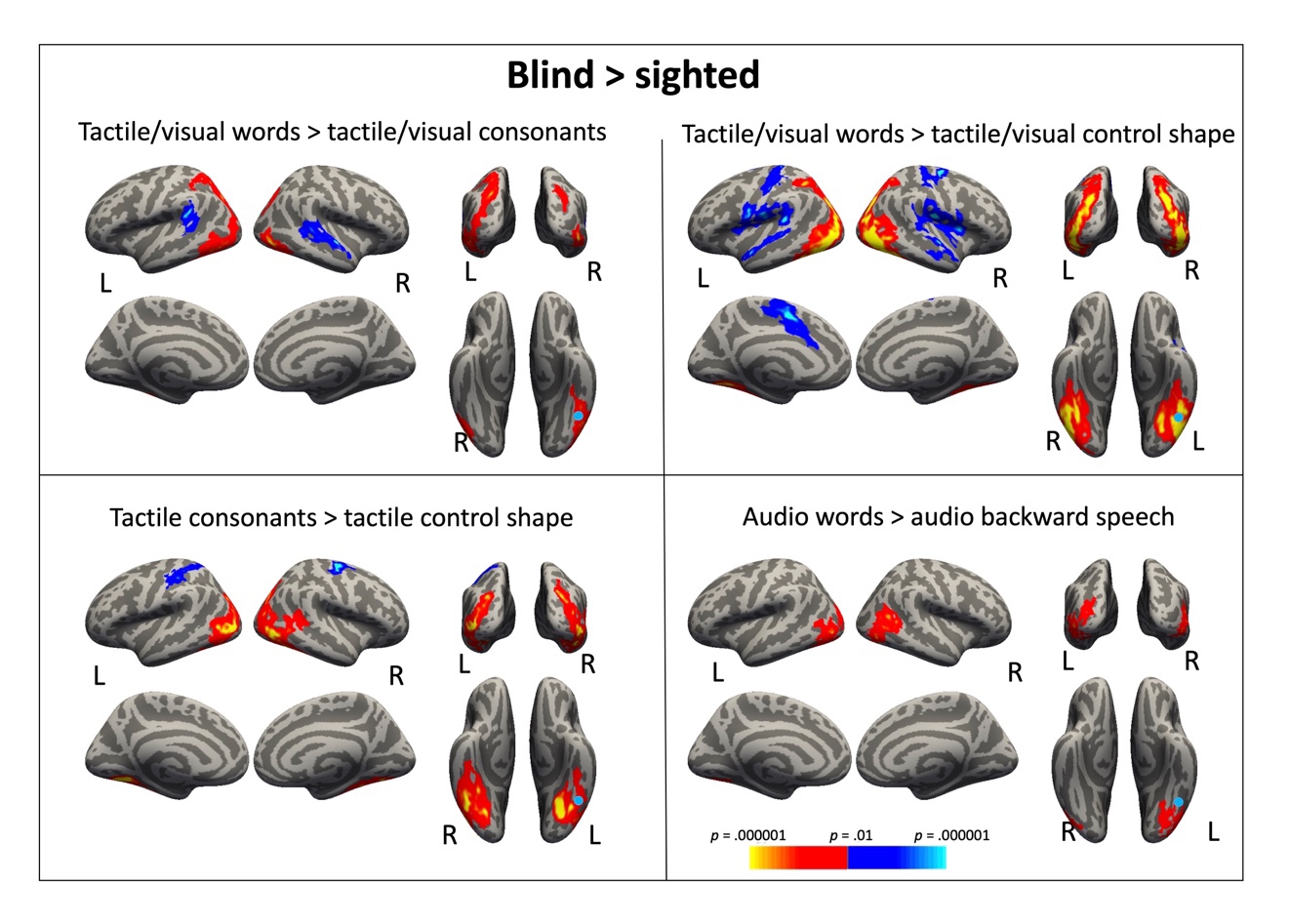

**Figure. S9a.** Between group difference in whole-cortex: blind > sighted (red), sighted > blind (blue), *p* < 0.05 cluster-corrected. Blue circles mark previously reported location of VWFA (MNI coordinate: -46, -53, -20). Note that this result should be interpreted with caution because visual and tactile reading tasks are inherently different.

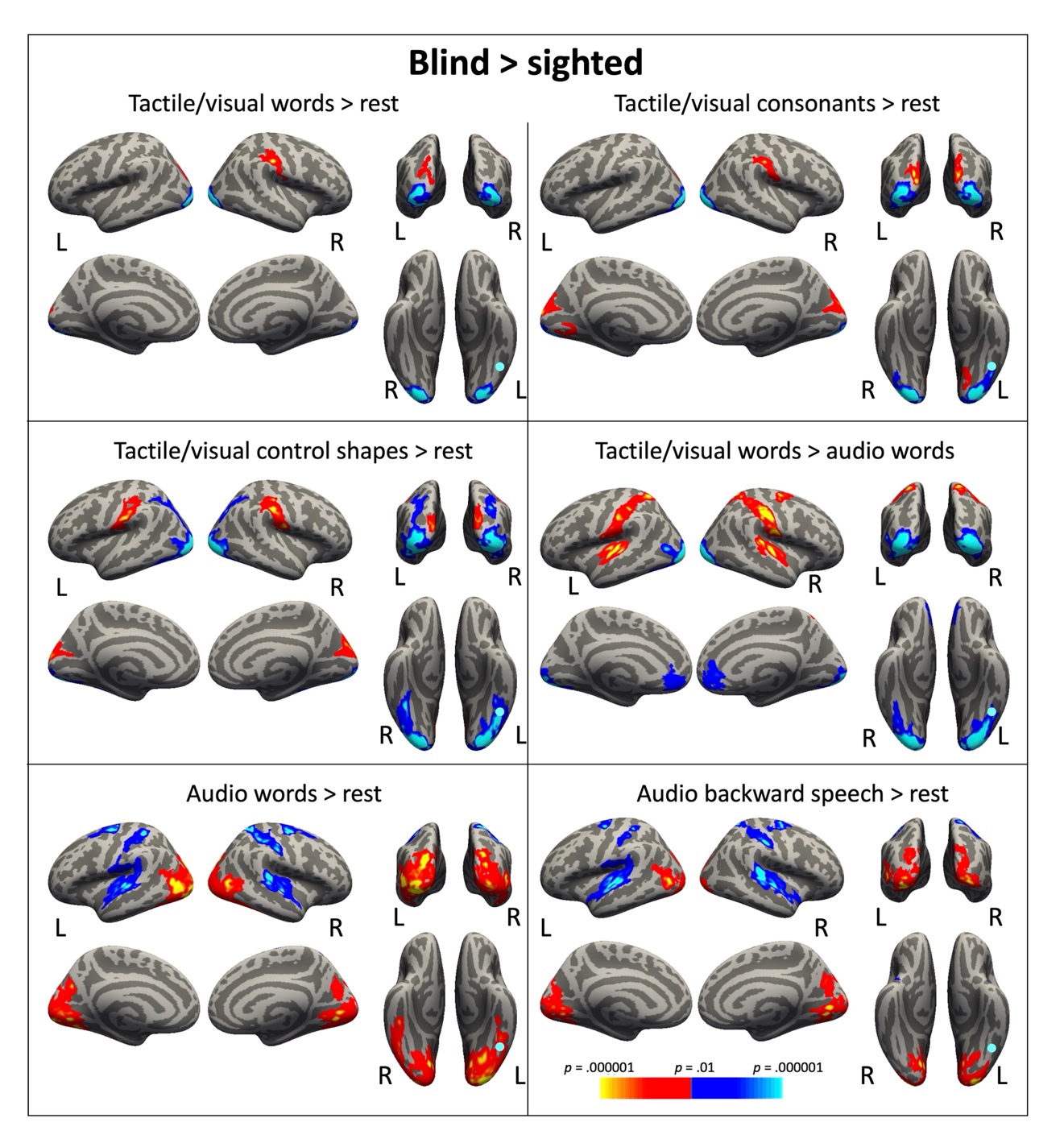

Figure. S9b. Between group difference in whole-cortex: blind > sighted (red), sighted > blind (blue), *p* < 0.05 cluster-corrected. Blue circles mark previously reported location of VWFA (MNI coordinate: -46, -53, -20). Note that this result should be interpreted with caution because visual and tactile reading tasks are inherently different.

**z**

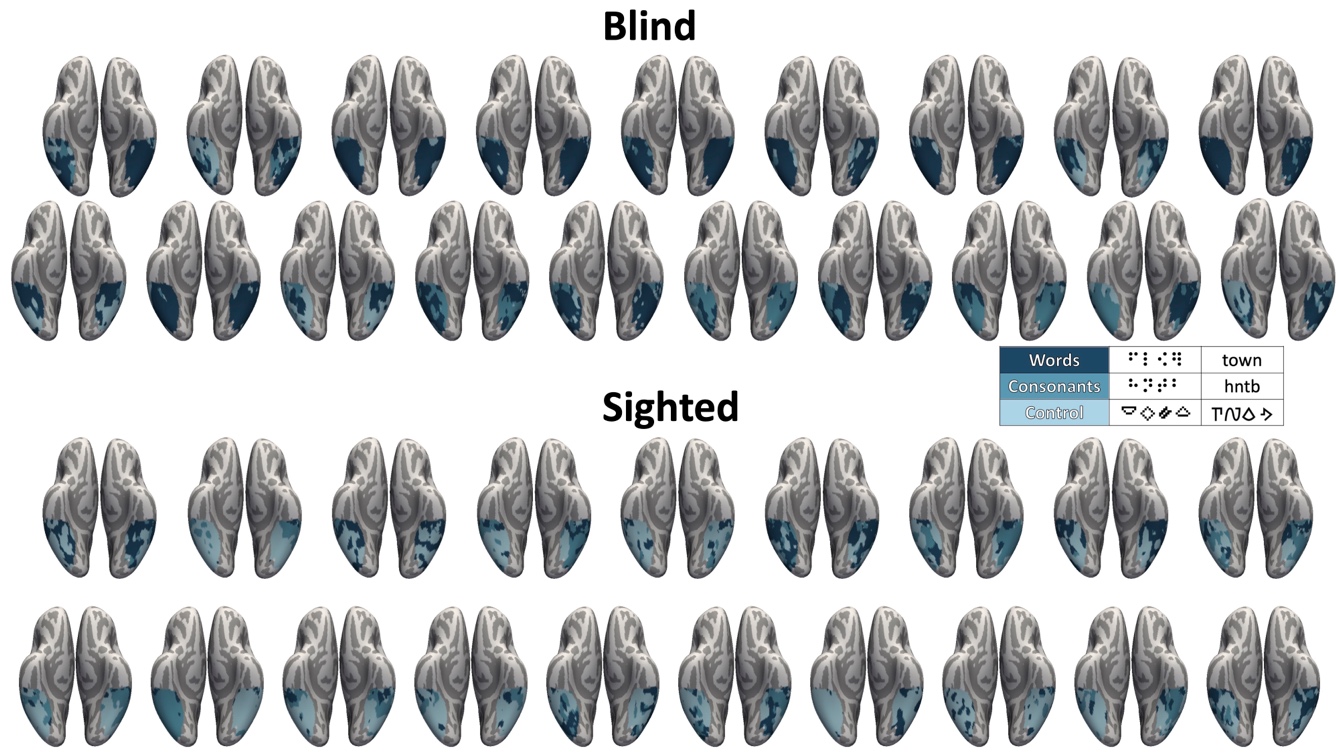

**Figure. S10a.** Winner-take-all map in vOTC for each participants during the reading task: words, consonant strings, control stimuli.

.

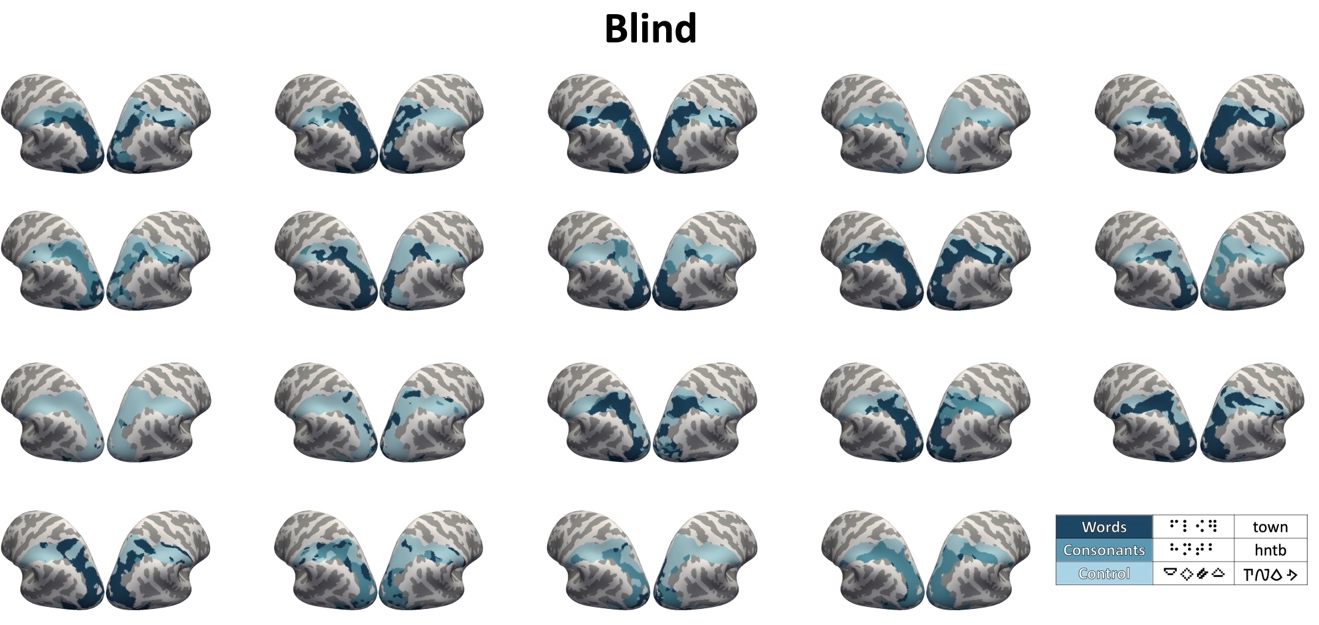

Figure. S10b. Winner-take-all map in posterior parietal cortex for each blind participants during the reading task: words, consonant strings, control stimuli.

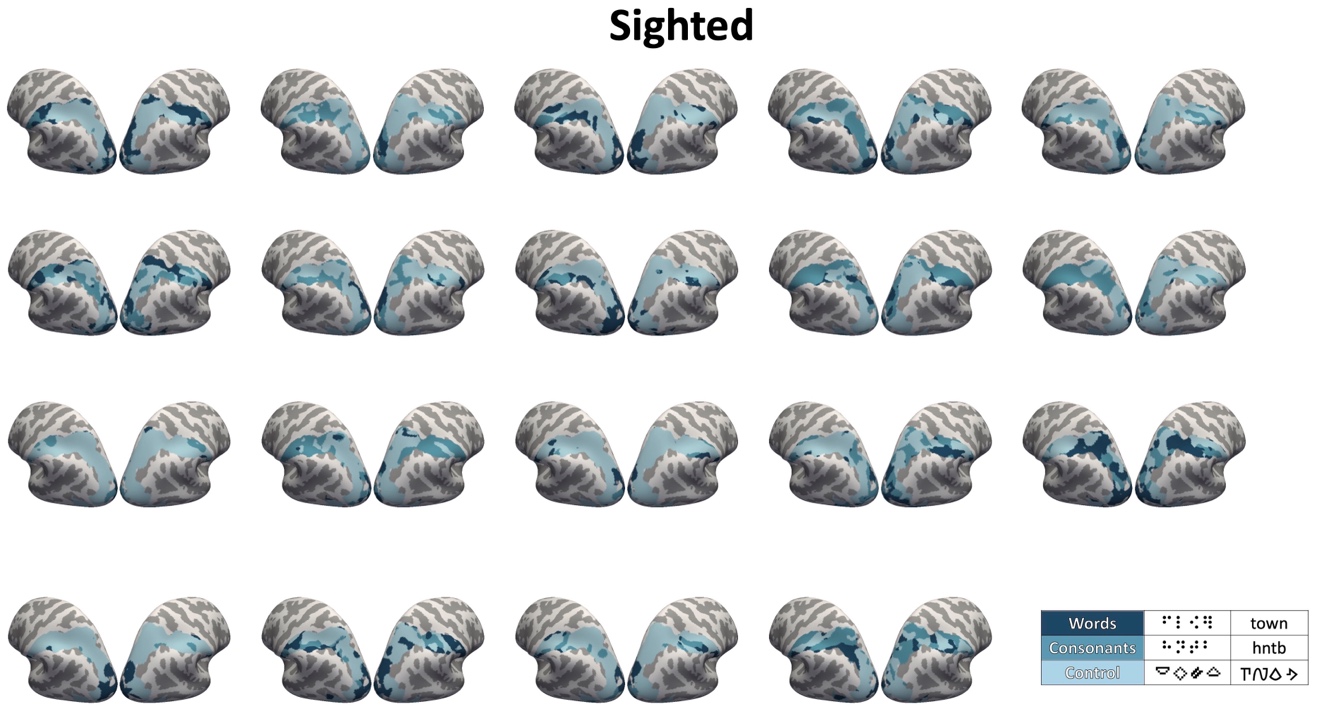

Figure. S10c. Winner-take-all map in posterior parietal cortex for each sighted participants during the reading task: words, consonant strings, control stimuli.

Table S1. Activated clusters in whole brain analysis

|  | | Brodmann’s areas | peak MNI coordinates | | | Peak z | Cluster size | | |
| --- | --- | --- | --- | --- | --- | --- | --- | --- | --- |
|  |  |  | X | Y | Z |  | vertices | | mm^2^ |
| **Blind** | |  | | | | | | | |
| **Braille words > rest** | |  | | | | | | | |
| Left hemisphere | Lateral/posterior occipital cortex (inferior occipital gyrus) | 19 | -42 | -73 | -4 | 5.79 | 4514 | 8935.79 | |
|  | Foveal confluence (middle occipital gyrus) | 18 | -26 | -92 | 14 | 4.97 |  |  | |
|  | Pericalcarine cortex/lingual gyrus | 17,18 | -6 | -72 | 4 | 4.74 |  |  | |
|  | Fusiform gyrus/inferior temporal gyrus | 37,21 | -40 | -57 | -13 | 5.26 |  |  | |
|  | Parieto-occipital cortex (middle occipital gyrus) | 18,19 | -20 | -85 | 21 | 5.28 |  |  | |
|  | Superior parietal lobule | 7 | -33 | -47 | 46 | 5.65 |  |  | |
|  | Supramarginal gyrus/postcentral gyrus | 7 | -42 | -41 | 39 | 4.25 |  |  | |
|  | Precentral gyrus | 6 | -52 | -2 | 43 | 4.8 | 782 | 1555.38 | |
|  | Inferior frontal gyrus (Pars opercularis) | 46 | -41 | 26 | 15 | 3.46 |  |  | |
|  | Middle frontal gyrus | 46 | -43 | 26 | 26 | 3.76 |  |  | |
| Right hemisphere | Superior parietal lobule | 7 | 20 | -60 | 60 | 5.38 | 4153 | 7924.29 | |
|  | Supramarginal gyrus/postcentral gyrus | 2,40 | 49 | -38 | 42 | 5.48 |  |  | |
|  | Parieto-occipital cortex (inferior parietal cortex/superior occipital gyurs/middle occipital gyrus) | 19,18 | 36 | -82 | 13 | 4.67 |  |  | |
|  | Fusiform gyrus/inferior temporal gyrus/middle temporal gyrus | 37 | 47 | -61 | -4 | 4.84 |  |  | |
|  | Pericalcarine cortex/lingual gyrus | 17,18 | 7 | -71 | 7 | 4.09 |  |  | |
|  | Precentral gyrus/ Inferior frontal gyrus (Pars opercularis) | 6,44 | -53 | -4 | 30 | 5.11 | 730 | 1482.12 | |
| **Auditory words > rest** | |  |  |  |  |  |  |  | |
| Left hemisphere | Superior temporal gyrus/transverse temporal gyrus/middle temporal gyrus | 38,39,22,42 | -47 | 0 | -18 | 6.02 | 4633 | 9736.91 | |
|  | Fusiform gyrus | 37 | -42 | -43 | -16 | 5.2 |  |  | |
|  | Lateral occipital cortex | 19,18 | -41 | -69 | -2 | 4.93 |  |  | |
|  | Lingual gyrus/cuneus cortex/pericalcarine cortex | 17,18 | -3 | -83 | 0 | 4.1 |  |  | |
|  | Inferior frontal gyrus (Pars opercularis)/middle frontal gyrus | 44, 6 | -52 | 14 | 14 | 4.65 | 313 | 619 | |
|  | Inferior frontal gyrus (Pars triangularis)/ lateral orbitofrontal gyrus/insula | 47, 45 | -39 | 31 | 0 | 4.45 |  |  | |
| Right  hemisphere | Superior temporal gyrus/transverse temporal gyrus | 38,22,42 | 49 | -14 | -5 | 6.39 | 4276 | 5221.69 | |
|  | Middle temporal gyrus | 39 | 51 | -59 | 5 | 4.38 |  |  | |
|  | Fusiform gyrus | 37 | 40 | -51 | -15 | 4.85 |  |  | |
|  | Lateral occipital cortex | 19 | 40 | -71 | 2 | 4.38 |  |  | |
|  | Pericalcarine cortex/lingual gyrus | 17,18 | 10 | -75 | 14 | 4.09 | 782 | 2036.1 | |
|  | Inferior frontal gyrus (pars triangularis)/insula | 45,46,47 | 43 | 30 | 0 | 4.86 | 1043 | 2025.2 | |
|  | Inferior frontal gyrus (Pars opercularis)/precentral gyrus | 44,45,6 | 51 | 12 | 16 | 3.82 |  |  | |
| **Braille words > audio words** | |  |  |  |  |  |  |  | |
| Left hemisphere | Superior parietal lobule/parieto-occipital cortex (superior occipital gyurs/middle occipital gyrus) | 19,18 | -22 | -77 | 29 | 5.67 | 2366 | 4095.12 | |
|  | Superior parietal lobule | 7 | -37 | -48 | 51 | 5.06 |  |  | |
|  | Supramarginal gyrus/postcentral gyrus | 2,40 | -39 | -32 | 35 | 4.61 |  |  | |
|  | Fusiform gyrus | 37 | -31 | -57 | -16 | 4.65 | 267 | 699.33 | |
|  | Lateral occipital cortex/inferior temporal gyrus | 37,19 | -44 | -67 | -5 | 4.04 | 383 | 667.97 | |
|  | Pericalcarine cortex/lingual gyrus/cuneus cortex | 17,18 | -15 | -93 | 0 | 4.51 | 348 | 754.36 | |
|  | Precentral gyrus/superior frontal gyrus | 4,6 | -31 | -11 | 50 | 4.17 | 192 | 429.25 | |
| Right hemisphere | Supramarginal gyrus/postcentral gyrus | 2,40 | 52 | -27 | 38 | 5.51 | 2702 | 4324.21 | |
|  | Parieto-occipital cortex (superior parietal lobule/superior occipital gyurs/middle occipital gyrus) | 19 | 24 | -78 | 38 | 5.07 |  |  | |
|  | Superior frontal gyrus | 6 | 10 | 1 | 51 | 4.99 | 683 | 1214.07 | |
|  | Precentral gyrus | 4 | 42 | -7 | 55 | 4.78 |  |  | |
| **Sighted** | |  |  |  |  |  |  |  | |
| **Visual words > rest** | |  |  |  |  |  |  |  | |
| Left hemisphere | Lingual gyrus/pericalcarine cortex (Foveal confluence/inferior occipital gyrus) | 17,18 | -9 | -94 | -8 | 6.66 | 2306 | 4896.03 | |
|  | Fusiform gyrus/inferior temporal gyrus | 37 | -35 | -45 | -20 | 6.32 |  |  | |
|  | Precentral gyrus | 4,6 | -52 | -5 | 46 | 4.99 | 1635 | 3230.98 | |
|  | Inferior frontal gyrus (parstriangularis/parsobitails) | 45,47,11 | -38 | 27 | 3 | 4.83 |  |  | |
|  | Inferior frontal gyrus (parsopercularis)/middle frontal gyrus | 44,46 | -47 | 13 | 23 | 4.83 |  |  | |
|  | Superior frontal cortex | 6,32 | -8 | 4 | 60 | 5.05 | 287 | 636.01 | |
|  | Inferior parietal lobule/superior parietal lobule | 7,19 | -26 | -64 | 27 | 4.91 | 849 | 1272.40 | |
|  | Middle temporal gyrus/superior temporal gyrus | 22 | -62 | -48 | -1 | 3.88 | 306 | 536.40 | |
| Right hemisphere | Calcarine/ Foveal confluence/lingual gyrus | 17,18 | 19 | -92 | -8 | 7.22 | 1604 | 3944.88 | |
|  | Fusiform gyrus/inferior temporal gyrus | 19,37 | 35 | -70 | -12 | 5.16 |  |  | |
|  | Superior parietal lobule | 7,19 | 31 | -59 | 44 | 4.86 | 389 | 591.7 | |
| **Audio words > rest** | |  |  |  |  |  |  |  | |
| Left hemisphere | Superior temporal gyrus/middle temporal gyrus | 22,42,38,39 | -60 | -23 | 1 | 6.59 | 2454 | 4822.44 | |
|  | Inferior frontal gyrus (parsopercularis)/middle frontal gyrus | 46,44 | -44 | 14 | 23 | 4.51 | 389 | 745.81 | |
|  | Inferior frontal gyrus (parstriangularis/parsobitails) | 45,47,11 | -36 | 29 | -3 | 4.19 |  |  | |
| Right hemisphere | Superior temporal gyrus/middle temporal gyrus | 22, 42, 38, | 59 | -30 | 8 | 6.18 | 1808 | 3638.92 | |
| **Visual words > audio words** | |  |  |  |  |  |  |  | |
| Left hemisphere | Lingual gyrus/ pericalcarine cortex (Foveal confluence/inferior occipital gyrus) | 17,18 | -20 | -88 | -8 | 6.91 | 2978 | 6164.4 | |
|  | Lateral occipital cortex/fusiform gyrus/interior temporal cortex | 19,37 | -35 | -45 | -20 | 5.23 |  |  | |
|  | Superior parietal lobule/inferior parietal lobule | 19,7 | -27 | -65 | 25 | 4.79 |  |  | |
| Right hemisphere | Lateral occipital cortex/ Pericalcarine cortex/ lingual gyrus | 17,18,19 | 19 | -92 | -8 | 6.1 | 1505 | 3662.98 | |
|  | Fusiform gyrus | 19,37 | 35 | -47 | -18 | 3.72 |  |  | |
|  | Superior parietal lobule | 7 | 28 | -56 | 45 | 4.14 | 569 | 1038.15 | |
